# A shared multi-feature population code for sensory reliability across mouse visual cortex

**DOI:** 10.64898/2026.05.21.726824

**Authors:** Elizabeth M. Edwards, Megan H. Lipton, Anne B. Sereno, Maria C. Dadarlat

## Abstract

Optimal use of sensory information requires the brain to represent uncertainty alongside sensory variables. Yet, the neural response properties that mediate reliability encoding remain undefined. Using large-scale two-photon calcium imaging of 140,000 neurons in primary visual cortex (V1) and higher visual areas (HVAs) during presentation of random dot kinematograms, we identified a common coding strategy in which sensory reliability is represented via coordinated changes in response gain, selectivity, and variability. These response properties are expressed at both single neuron and population levels and are consistent across areas, with quantitative differences in magnitude that track retinotopic drive and functional specialization. Together, our results establish reliability encoding as a general cortical computation implemented through a shared population code, providing empirical constraints on probabilistic theories of perception and hierarchical models of visual processing.

## INTRODUCTION

Accurate perception requires the brain to resolve ambiguity by integrating noisy, incomplete, or conflicting sensory inputs into coherent representations that support adaptive behavior. Accumulating evidence suggests that the brain encodes not only stimulus features but also their reliability, thereby forming an internal statistical model of the world that supports inference under uncertainty (1, 2, 3, 4, 5). A prominent theory posits that uncertainty is multiplexed with stimulus content in sensory cortical populations, such that the neurons encoding a particular world state variable also carry information about the reliability of that estimate (1, 4, 6, 7). Within this framework, perceptual decisions are based on probability distributions over task-relevant latent variables (e.g., the direction of motion of a visual stimulus) rather than point estimates. Human and animal behavior supports such probabilistic accounts of perception (8, 9, 10); yet, the neural code by which uncertainty about sensory variables is represented remains incompletely resolved.

In the visual system, where stimulus uncertainty can be precisely manipulated and mapped onto well-characterized neural population responses, the question of how neurons encode sensory reliability becomes tractable. Evidence from both ends of the visual hierarchy suggests that reliability is encoded at multiple levels of processing. Population-level signals in early visual cortex systematically covary with stimulus reliability (6), while single neurons in higher-order parietal and prefrontal areas represent confidence in perceptual decisions (11, 12, 13). However, these observations have primarily been made at different levels of description (single neurons vs. populations) and in different parts of the visual processing hierarchy. Thus, it remains unclear how reliability information is processed and refined across sensory cortex in the response properties of single neurons and neural populations.

A central challenge in addressing reliability encoding is to dissociate changes in the reliability of a sensory signal from changes in the low-level stimulus properties that independently drive neural responses. For example, reducing the contrast of oriented grating stimuli simultaneously alters effective stimulus drive alongside reliability, making it difficult to attribute changes in neural activity specifically to reliability (6, 14, 15). Random dot kinematograms (RDKs) provide a principled solution to this challenge by manipulating the ratio of signal-to-noise dots (motion coherence) to smoothly degrade the reliability of the sensory feature (motion direction), while preserving low-level stimulus properties (dot contrast, density, spatial statistics; (16, 17, 18, 19, 20)). Changing motion coherence modulates the reliability of the directional signal in an ethologically-relevant way, with low-coherence RDKs consistently driving slower, less-accurate behavior (18, 19, 20, 21, 22, 23, 24). Importantly, 0% coherence RDKs also conserve total motion energy but lack a net directional signal, providing an empirical baseline for visual drive and a mechanism for within-experiment dissociation of reliability-dependent modulation from coherence-independent visual responses. Thus, RDKs are well-suited for isolating the neural code for sensory reliability.

In primate middle temporal (MT) cortex, coherence modulates single neuron response amplitudes, with more reliable motion signals driving stronger direction-selective responses (16, 25, 26, 27, 28, 29, 30). Similar coherence-dependent response scaling occurs across mouse primary visual cortex (V1) and functionally-specialized higher visual areas (HVAs; (19, 31, 32)), establishing the mouse visual system as a convenient model for studying reliability encoding across a well-characterized cortical hierarchy (33, 34, 35, 36). This conserved relationship between coherence and response magnitude implicates gain modulation as one component of the reliability code. However, whether gain modulation alone fully accounts for how reliability is represented at the level of single neurons and populations remains unresolved.

Two additional response properties have been proposed as candidates for encoding reliability alongside gain modulation. First, tuning curve sharpening, in which more reliable stimuli increase directional selectivity (7, 37, 38, 39). Second, reduced trial-to-trial variability, in which more reliable stimuli drive more consistent responses (40, 41). Prior studies have typically examined these mechanisms in isolation and within single brain areas, leaving open whether they act independently or in concert across single neurons and populations to shape reliability signals.

Here, we test whether single neurons and neural populations share a common set of response properties for encoding sensory reliability, and whether those properties are conserved or scaled across the visual hierarchy. To address these questions, we used large-scale two-photon calcium imaging to record the activity of ∼140,000 neurons across mouse V1 and six HVAs during passive viewing of behaviorally-calibrated coherent motion. We show that single neurons encode changes in stimulus reliability through coordinated modulation of response gain, tuning width, and variability, rather than via isolated changes in any single response property. This multi-feature reliability coding scheme is preserved at the population level across V1 and HVAs, though with quantitative differences in expression reflecting functional specialization and retinotopic drive from the stimulus. Together, these results reveal a unified reliability code in mouse visual cortex that is shared across single neurons and neural populations and scaled across visual areas.

## RESULTS

### Behavioral sensitivity to motion coherence defines reliability regimes

An animal’s ability to infer visual motion direction from a random dot kinematogram (RDK) provides a behavioral estimate of the reliability of the visual signal. To characterize this relationship, we trained mice on a two-alternative forced-choice (2AFC) task in which they rotated a wheel to report the horizontal flow direction of an RDK displayed within the binocular visual field (Fig. 1A-B; (42, 43)). Choice accuracy increased systematically with motion coherence, and mice achieved a maximum performance of 87% ± 3% correct choices at 100% coherence (mean ± SEM; *n* = 5 mice; Fig. 1C). Fitting a mean psychometric function yielded a coherence threshold of 33% for 70% correct performance, consistent with previous work using similar stimuli (19, 20).

**Figure 1:**
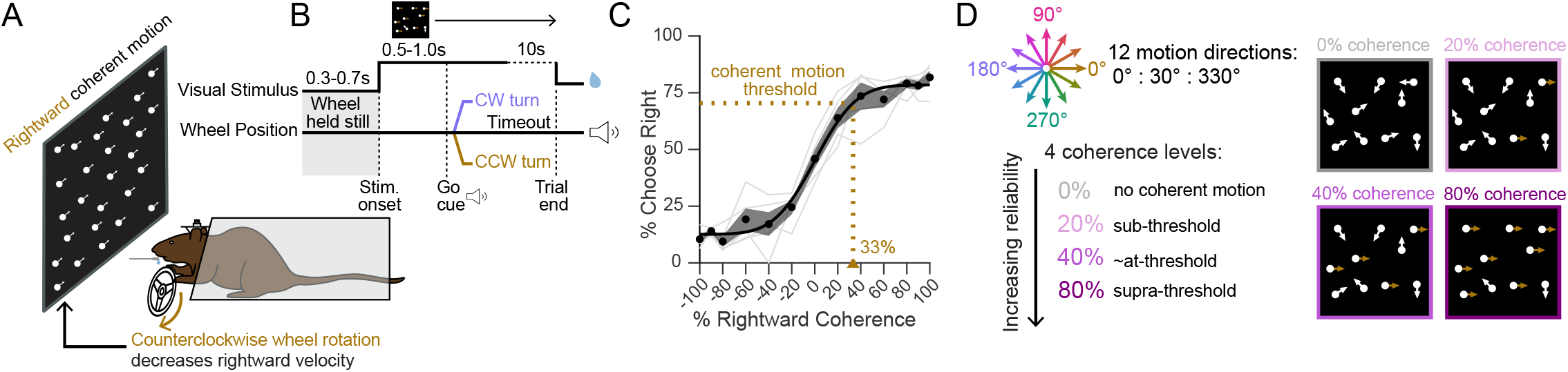
Motion coherence provides a quantifiable proxy for sensory reliability. **(A)** Schematic of two-alternative forced choice task. Head-fixed mice reported the horizontal motion direction (left or right) of a bilaterally presented random dot kinematogram (RDK) by rotating a wheel with their forepaws. **(B)** Trial structure. Mice initiated trials by holding the wheel still. The stimulus then appeared but visual flow speed remained fixed and reward unavailable until an auditory go cue played. Correct wheel movements were rewarded, while incorrect or missed responses (>10s) were punished with a white noise burst. CW, clockwise; CCW, counterclockwise. **(C)** Psychometric performance of expert mice (*n=5*). Circles indicate group means; black line, psychometric fit; gray lines, individual mice; shading, SEM. Leftward stimuli are represented with negative coherence. The threshold for 70% correct performance is indicated. **(D)** Schematic of RDK stimuli used during passive viewing. Stimuli were presented at four coherence levels (0, 20, 40, 80%) in 12 equally-spaced motion directions (0° – 330°, colored arrows), selected according to behavioral sensitivity to coherence measured in (C).

We used this measurement of behavioral sensitivity to define coherence levels for subsequent passive neural recordings, selecting values that spanned a broad range of perceptual difficulty: 0% (no coherent motion), 20% (below behavioral threshold), 40% (near threshold), and 80% (above threshold). For each non-zero coherence, stimuli were presented in 12 evenly-spaced motion directions (0° – 330°; Fig. 1D).

### Large-scale multi-area recordings reveal distributed single-neuron representations of coherent motion

The stimulus set defined in Fig. 1D was passively-viewed by awake, head-fixed mice expressing the calcium indicator GCaMP6s in excitatory neurons (TRE-GCaMP6s::CaMKII-tTA; Fig. 2A). Screen location and stimulus parameters were identical to those used during behavior. We recorded neural activity via single-plane, two-photon calcium imaging over the right visual cortex. In total, we recorded approximately 138,000 neurons across eight mice (Fig. 2B; 3,292 ± 74 recorded simultaneously per session; 42 sessions; Table S1). Stimulus-evoked responses were taken as deconvolved fluorescent traces, averaged over the two-second stimulus presentation window (44, 45).

**Figure 2:**
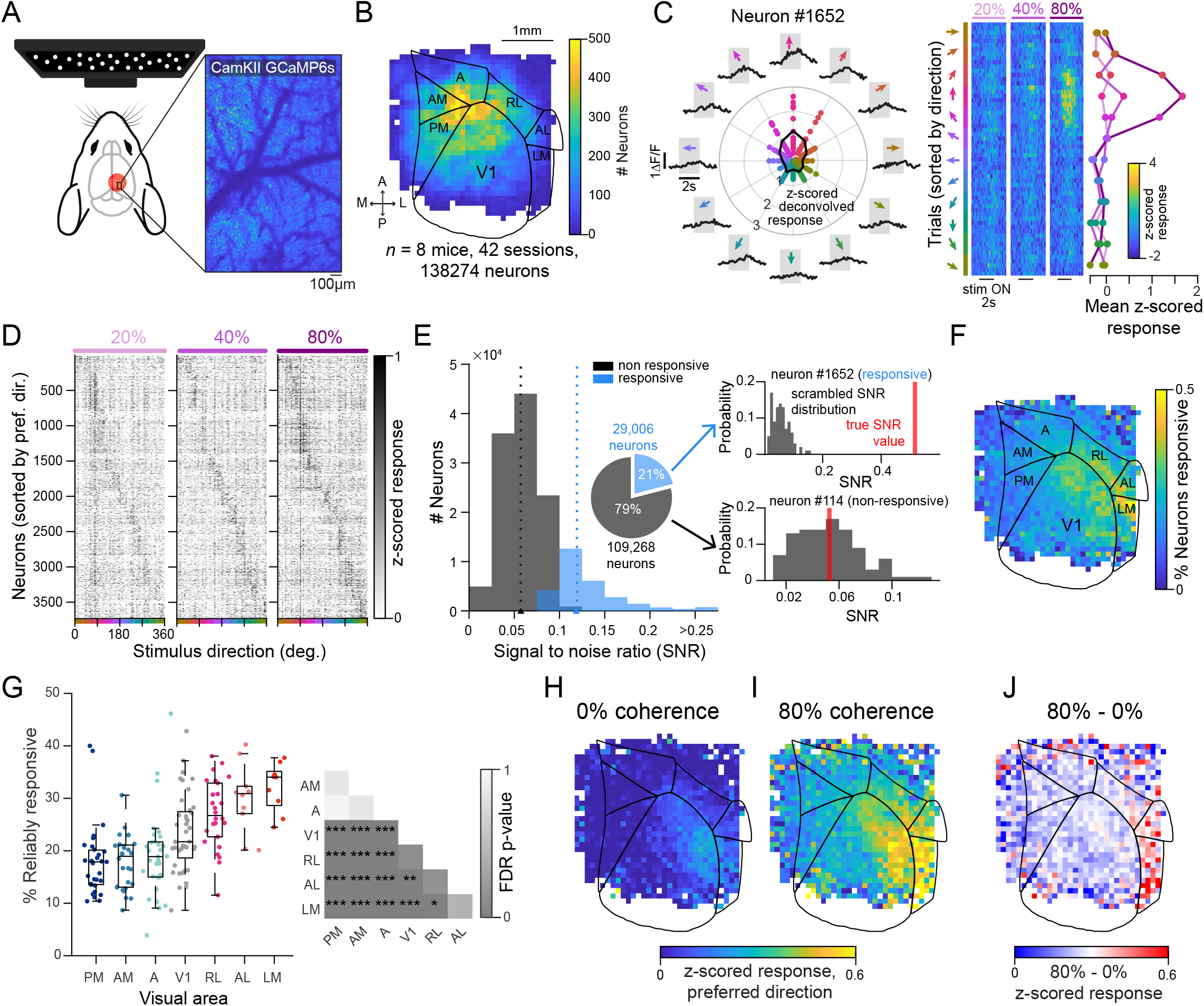
Large-scale two-photon calcium imaging reveals lateralized responses to coherent motion. **(A)** Experimental setup. Neuronal activity in right visual cortex was recorded using two-photon calcium imaging while mice passively viewed random dot kinematograms (RDKs) varying in motion direction (0° – 330°) and coherence (0, 20, 40, 80%). Right, example GCaMP6s maximum projection image. **(B)** Spatial distribution of all recorded neurons registered to the Allen Institute Mouse Common Cortical Framework (CCF; *n* = 8 mice, 42 sessions, 138,274 neurons). Grid size: 0.1mm x 0.1mm. Scale bar = 1mm. **(C)** Directionally-selective responses of an example neuron. Left, time-averaged deconvolved responses to each motion direction (colored dots) with fitted tuning curve (black line); outer ring shows normalized trial-averaged fluorescence traces (ΔF/F). Gray shading indicates 2s stimulus window. Right, single-trial activity sorted by direction and coherence (heatmaps) with trial-averaged responses as a function of direction for each coherence level (pink lines). **(D)** Population responses from a single session (*n* = 3,756 neurons), sorted by preferred direction across stimulus conditions. **(E)** Neuronal responsiveness classification by signal-to-noise ratio (SNR). Left, SNR distribution; responsive neurons in blue. Right, example true vs. shuffled SNR comparisons used to threshold responsiveness. **(F)** Spatial distribution of responsive neurons overlaid with the Allen CCF. **(G)** Fraction of responsive neurons by visual area. Points indicate session-level estimates. Right, false discovery rate (FDR)-corrected *P* values from pairwise linear mixed effects model (LMM) contrasts. **(H-J)** Spatial distribution of trial-averaged responses at 0% coherence (H), 80% coherence in each neuron’s preferred direction (I), and their difference (J), highlighting enhanced sensitivity to coherent motion in lateral visual areas. ^†^*P <* 0.1, \**P* < 0.05, \*\**P* < 0.01, \*\*\**P* < 0.001. See also Figs. S1 and S2.

To characterize responses by cortical location, fields-of-view from each imaging session were registered to the Allen Institute Mouse Common Cortical Framework (CCF) using surface vasculature patterns and retinotopic mapping (Figs. 2B and S1; (33, 45, 46, 47)). Neurons were recorded across primary visual cortex (V1) and six adjacent higher visual areas (HVAs; PM = posteromedial, AM = anteromedial, A = anterior, RL = rostrolateral, AL = anterolateral, and LM = lateral medial; Table S2).

Consistent with previous reports (19, 32, 31), individual neurons exhibited directionally-specific responses to RDK motion that increased with coherence (Fig. 2C). For each neuron, the preferred motion direction was estimated by fitting sine and cosine basis functions to responses across all non-zero coherence stimuli (45).

Despite this selectivity, neurons responded variably across trials, even to their preferred motion directions (Fig. 2D). Neuronal responsiveness was therefore quantified using the signal-to-noise ratio (SNR) of the fitted tuning function relative to a null-SNR bootstrapped distribution derived from permuted responses (Figs. 2E and S2; (45)). With this approach, 21% of neurons (29,006) were classified as reliably responsive (median SNR: 0.12, interquartile range (IQR): 0.11 to 0.14; Table S1). Subsequent analyses were restricted to this reliably responsive subpopulation.

Responsive neurons were distributed across visual areas, with the highest concentration in anterior and lateral V1 and the adjacent lateral HVAs: RL, AL, and LM (Figs. 2F and S2; Table S2). Across sessions, the fraction of responsive neurons was significantly higher in AL and LM compared to medial visual areas (PM, AM, A; Figs. 2G and S2B). This lateralized activation pattern is consistent with the central visual field placement of the stimulus (Fig. 2A) and prior reports of central visual field dominance in mouse lateral visual cortex (31, 33, 48, 49).

To determine whether the enhanced lateral responsiveness specifically reflected sensitivity to coherent motion or simply stronger visual drive from the centrally-placed stimulus, we compared responses at 0% motion coherence, which contains motion energy but lacks a coherent direction, to responses at 80% motion coherence in each neuron’s preferred direction. Responses at 0% coherence were modestly elevated in lateral V1 and lateral HVAs (Fig. 2H), but became further amplified at 80% coherence (Fig. 2I-J), confirming selective sensitivity to directional motion in lateral areas.

### Single neurons encode sensory reliability through coordinated modulation of response gain, tuning width, and variability

Having identified neurons that reliably respond to coherent motion (Fig. 2), we next asked whether changing coherence modulated their responses in a manner consistent with the three previously-presented candidate encoding mechanisms (Fig. 3A): changes in response gain, tuning width, or trial-to-trial response variability. Because these features cannot be estimated for single neurons on individual trials, we quantified each mechanism using tuning curves computed across repeated stimulus presentations (Fig. 3B,C).

**Figure 3:**
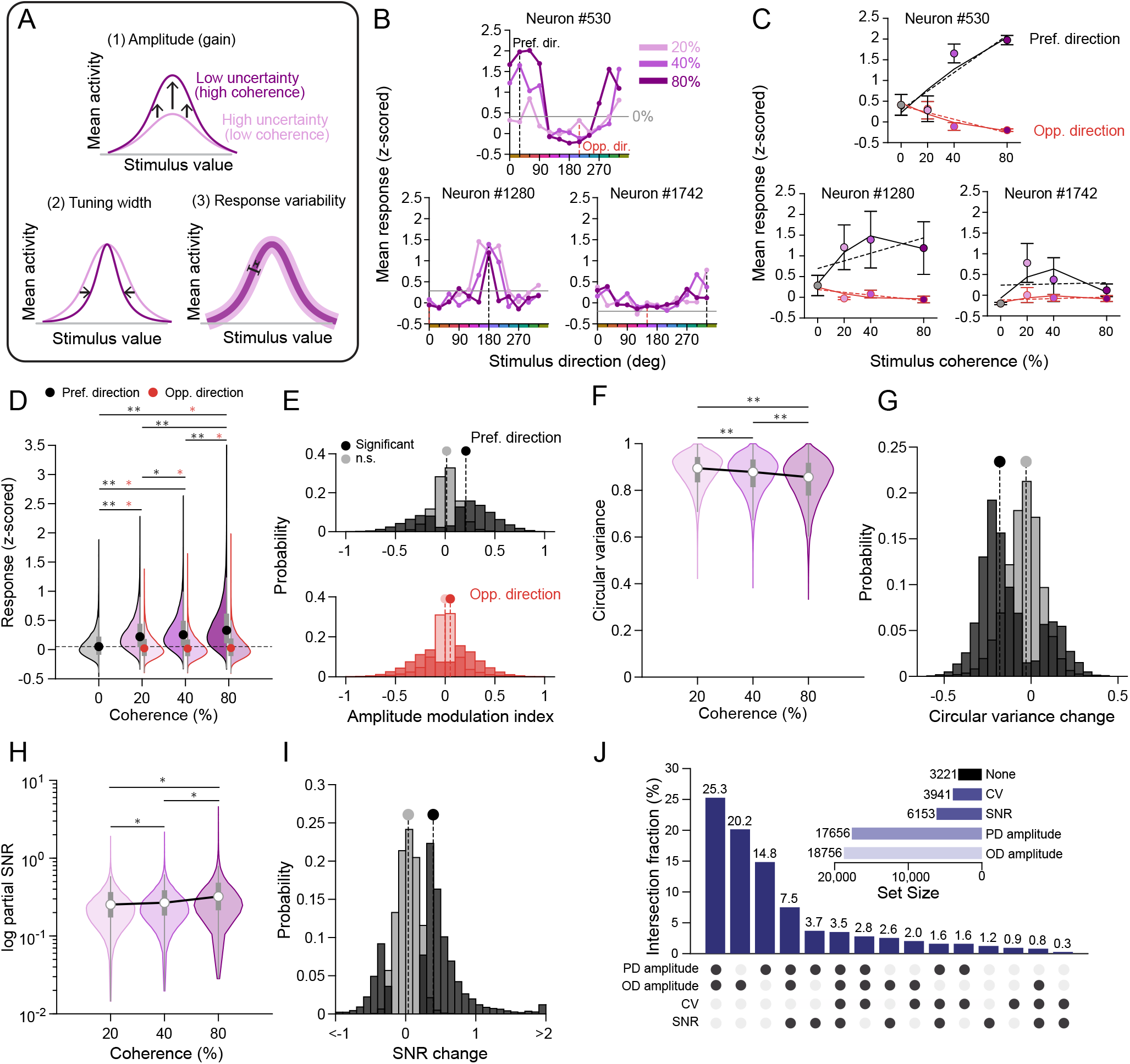
Single neurons use a structured, multi-feature encoding scheme for sensory reliability. **(A)** Schematic illustrating three candidate mechanisms by which increasing sensory reliability (motion coherence) may modulate single-neuron directional tuning: (1) increased response amplitude, (2) tuning width narrowing, (3) reduced trial-to-trial variability. **(B)** Trial-averaged responses of neurons as a function of coherence (pink color) exhibiting the three candidate mechanisms outlined in (A). Vertical black dashed line, preferred stimulus direction; vertical red dashed line, opposite direction (180° offset). Horizontal gray line, average response to 0% coherence. **(C)** Response functions for neurons in (B), showing mean response as a function of coherence. Black outline, preferred direction response; red outline, opposite direction response. Error bars indicate standard error of the mean response. Curves are best fitting linear (dashed) and quadratic (solid) lines. **(D)** Population distribution of trial-averaged z-scored deconvolved activity for responsive neurons in their preferred (black outline) and opposite (red outline) directions as a function of stimulus coherence. Circles indicate medians; gray bars denote interquartile range and whiskers. **(E)** Distribution of single-neuron amplitude modulation indices (AMI) for preferred direction (top) and opposite-direction (bottom) responses. Dark shading indicates neurons exhibiting significant modulation relative to shuffled controls. **(F)** Circular variance (CV) of responsive neurons as a function of coherence. **(G)** Distribution of single-neuron coherence-dependent changes in CV from 80% to 20% coherence, shown as in (E). **(H)** Partial signal to noise ratio (SNR) as a function of coherence (logarithmic y-axis for visualization). **(I)** Distributions of single-neuron coherence-dependent changes in partial SNR, shown as in (E). Values binned to [-1,2] for visualization. **(J)** Upset plot showing the fraction of responsive neurons (*n* = 29,006) exhibiting each combination of modulation mechanisms. Dots indicate the mechanism set corresponding to each bar. Inset, total significant neuron counts for each mechanism. Statistics in (D), (F), and (H) indicate pairwise comparisons between coherence levels using two-sided Wilcoxon sign-rank tests on mouse averages, FDR-corrected for multiple comparisons (*n* = 8 mice). \**P* < 0.05, \*\**P* < 0.01

We first examined gain modulation, a mechanism by which changes in stimulus reliability selectively rescale neural response amplitudes to enhance the true stimulus response. Consistent with previous work (18, 19, 32), responses in each neuron’s preferred direction tended to increase monotonically with coherence (Fig. 3D), while responses in the opposite direction (180° offset) were typically suppressed relative to 0% coherence. To quantify these changes, we computed an amplitude modulation index for responses in both the preferred and opposite directions for each neuron: *AMI* = (*R*_80%_ −*R*_20%_)/(*R*_80%_+ *R*_20%_) (Fig. 3E). A neuron was classified as *significantly* modulated if its true AMI exceeded the 90th percentile of a shuffled response distribution. Preferred-direction (PD) AMI values were strongly right-skewed (median: 0.21, IQR: -0.18 to 0.38), with 60.8% of responsive neurons (*n* = 17,656) showing significant coherence-dependent amplitude changes (Fig. 3E, top). In contrast, opposite-direction (OD) AMI values were approximately zero-centered (median: 0.05, IQR: -0.22 to 0.23), with 64.6% of responsive neurons (*n* = 18,756) modulated, indicating that opposite-direction responses are not uniformly suppressed by increasing coherence (Fig. 3E, bottom).

We next examined tuning width modulation, which considers how a neuron’s directional specificity changes with coherence. To estimate tuning width, we calculated the circular variance (CV) of each tuning curve, a shape-agnostic measure of directional selectivity where lower values indicate narrower tuning (50). CV consistently decreased with coherence (Fig. 3F). Because CV is bounded between 0 and 1, we quantified modulation magnitude as the difference in CV between high (80%) and low (20%) coherence. Significance was again computed relative to a shuffled null distribution. Neurons with significant CV modulation (*n* = 3,941) showed a strongly left-skewed distribution of CV changes (median: -0.18, IQR: -0.25 to -0.05; Fig. 3G), indicating more directionally-precise single neuron responses when signal reliability (motion coherence) increases. Although fewer neurons reached significance for tuning width changes compared to gain modulation, the population-level distribution was consistently left-skewed (median: -0.04; Fig. 3G), indicating that coherence-dependent sharpening is also a broadly expressed feature of the reliability code.

Finally, we considered single neuron trial-to-trial response variability, which should decrease with stronger stimuli (40, 41). We estimated variability by computing the SNR separately for each coherence level (45), a value we term the *partial* SNR. Partial SNR increased with coherence (Fig. 3H), indicating more consistent responses across trials when reliability is stronger. Significant partial SNR changes (*n* = 6,153 neurons), again computed by shuffling, were strongly right skewed (median = 0.39, IQR: 0.27 to 0.56; Fig. 3I), indicating that reduced response variability is a third feature of the reliability code.

Given all three modulation mechanisms were broadly present in cortical responses, we then asked whether coherence modulates these features independently or drives coordinated modulation across the three features. Intersection analyses revealed that amplitude modulation was the most common pattern, with 25.3% of responsive neurons showing significant amplitude changes in their preferred and opposite directions simultaneously (Fig. 3J). When neurons exhibited CV or SNR changes, these effects almost always co-occurred with gain modulation (≤ 2% of CV or SNR changes occurred without accompanying PD or OD gain modulation). Together, these results indicate that visual motion reliability is encoded through a robust, multi-feature modulation of gain, tuning width, and variability, with gain serving as the dominant component of this code.

### Reliability-encoding mechanisms are shared across visual cortical areas but differ in magnitude

To this point, we have broadly considered single-neuron encoding mechanisms, regardless of the spatial location of recorded neurons. However, functional and retinotopic differences across mouse HVAs raise the possibility that reliability encoding could vary across visual areas in mechanism prevalence or magnitude (32, 33, 34, 36, 51). It is possible, therefore, that area-specific biases for certain mechanisms could explain the comparatively lower prevalence of CV and SNR observed in Fig. 3J. To examine this, we compared the prevalence and magnitude of the reliability-encoding mechanisms across V1 and six HVAs.

Contrary to this specialization hypothesis, visual areas employed reliability-encoding mechanisms at broadly similar relative rates (Fig. 4A). Across areas, gain modulation was substantially more prevalent than CV or SNR modulation, consistent with its role as the dominant component of the reliability code (generalized linear mixed effects model (GLMM); odds ratio relative to PD AMI: CV, 0.10 [95% CI, 0.10-0.11]; SNR, 0.17 [0.17-0.18]). However, mechanism prevalence showed modest area dependence: lateral visual areas (RL, AL, LM) had higher probabilities of expressing gain and SNR modulation relative to medial areas (A, AM, PM; GLMM pairwise contrasts, Fig. S3).

**Figure 4:**
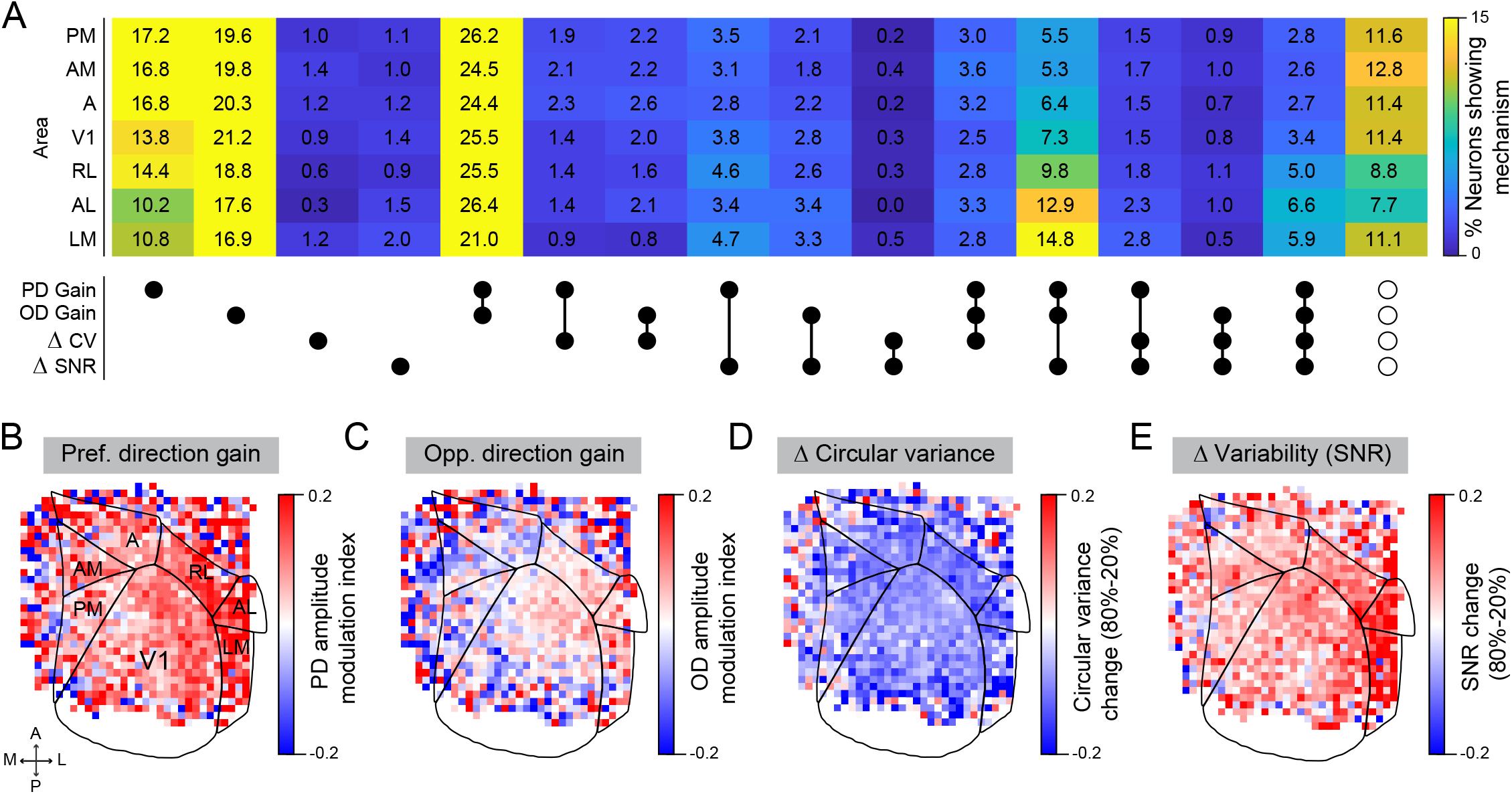
A shared set of reliability-encoding mechanisms is expressed across visual cortex with area-specific magnitudes. **(A)** Fraction of responsive neurons within each visual area that exhibit distinct combinations of reliability-encoding mechanisms. Rows correspond to visual areas; dots indicate the mechanism set associated with each bar. **(B)** Spatial distribution of preferred direction (PD) gain, quantified as amplitude modulation index (AMI). Area boundaries from the Allen Institute Mouse CCF are overlaid (*n* = 8 animals; 29,006 responsive neurons). Grid size: 0.1mm x 0.1mm. **(C-E)** Same as (B) for opposite direction gain (OD; C), circular variance change (D) and SNR change (E). See also Fig. S3.

We then evaluated the spatial distribution of encoding strength across visual cortex to determine whether mechanism magnitude was uniform within each area (Fig. 4B-E). Strong coherence-dependent increases in gain (PD and OD) and SNR formed a distinct lateral band, aligning with regions receiving the strongest visual drive from the centrally positioned visual display (Fig. 4B-C, E). In contrast, circular variance was more hierarchically distributed, with moderately larger decreases in HVAs compared to V1 (Fig. 4D).

### Single-trial population-level encoding of sensory reliability employs the same mechanisms as single neurons

So far, we have shown that sensory reliability shapes the responses of single neurons to coherent visual motion across trials. However, at the timescale of a single trial, reliability must be encoded in the distribution of responses across a neural population (1, 7, 52). Having shown that single neurons exhibit coordinated reliability-dependent modulation of response amplitude, tuning width, and response variability (Fig. 3), we next asked whether the same mechanisms are expressed in single-trial population responses. To address this, we constructed population tuning curves for each visual area by averaging preferred-direction-aligned responses across neurons on each trial (Figs. 5A and S4A), preserving single-trial variability while capturing the population response structure. For each population tuning curve, we computed the same four tuning features as in the single-neuron analyses: tuning curve amplitude (preferred and opposite directions), circular variance, and trial-to-trial response variability. For the latter, we used a bounded [0,1] population SNR metric (pSNR) quantifying alignment of each trial’s population response to a session-averaged template (see Methods).

**Figure 5:**
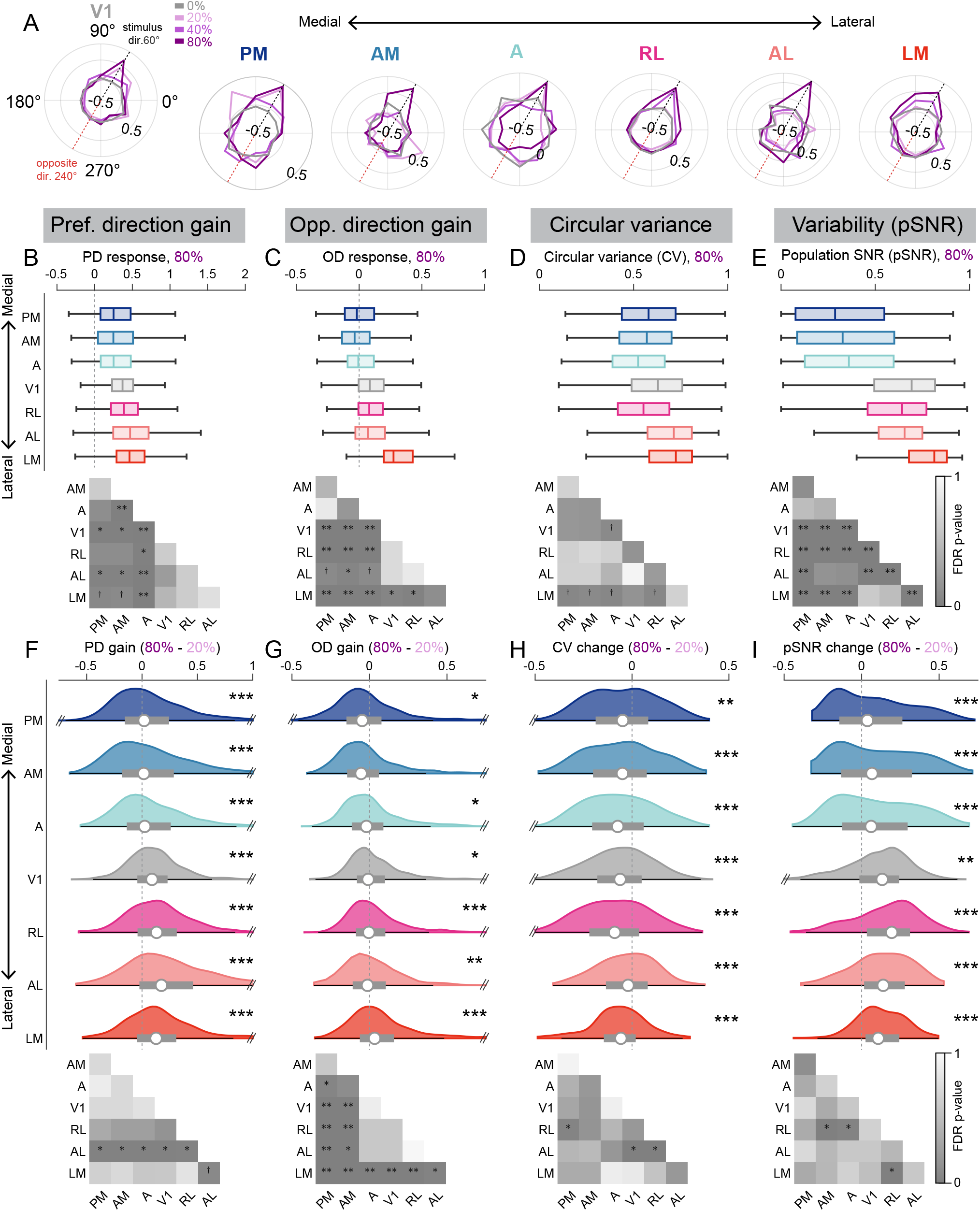
Population-level responses maintain single-neuron reliability-encoding mechanisms. **(A)** Example single-trial population tuning curves for each visual area. Responses shown for example 60° direction trials across coherence levels (pink shades) and mean 0% coherence response (gray). **(B)** Preferred-direction (PD) population tuning curve response amplitude at 80% coherence. Bottom, pairwise comparisons between visual areas. **(C-E)** Same as (B) for opposite direction (OD) response amplitude (C), circular variance (D), and population bounded SNR (pSNR; E). **(F)** Distribution of change in PD amplitude from 80% to 20% coherence. Circles, medians; gray bars, interquartile ranges. Dashed vertical line, zero change. Statistics indicate comparison of distribution to zero for each area. Truncated for visualization. **(G-I)** Same as (F) for OD amplitude (G), circular variance (H), and pSNR (I). Bottom, pairwise comparisons of modulation across visual areas. All statistics indicate bootstrap LMM contrasts with FDR correction. ^†^*P <* 0.1, \**P* < 0.05, \*\**P* < 0.01, \*\*\**P* < 0.001. See also Figs. S4 and S5.

At 80% coherence, population encoding of visual motion was stronger in V1 and lateral areas (RL, AL, LM) compared to medial areas (PM, AM, A), instantiated as higher PD response amplitude and pSNR (Fig. 5B-C,E). Population circular variance showed a similar but weaker trend, with marginally higher values in LM than other areas (Fig. 5D). This lateral dominance of sensory encoding was similarly visible at 20% coherence (Fig. S4B-E) and mirrors single-neuron patterns (Fig. S5A-D), consistent with stronger retinotopic drive to lateral visual areas from the centrally-positioned stimulus.

Despite baseline differences in encoding strength, coherence-dependent modulations were consistently implemented across visual areas (Fig. 5F-I). Across areas PD amplitude and pSNR increased with coherence, while circular variance decreased (Figs. 5F,H-I and S4F,H-I). The one exception to this consistent pattern was OD amplitude, which significantly decreased in V1 and medial areas (PM, A), but not in lateral areas (LM, AL, RL), suggesting that lateral populations respond with less directional specificity at high coherence (Figs. 5G and S4G). Together, these results provide evidence that reliability encoding across mouse visual cortex is not area-specific, but instead reflects a shared, multi-feature computational strategy that is functionally scaled across the visual hierarchy.

### Single-trial population-level decoding supports a global reliability code across visual cortex

Having characterized how reliability is encoded in population activity across visual areas, we next asked how the observed quantitative differences in encoding strength relate to differences in directional information available in each population. We decoded motion direction from single-trial population activity using a linear decoder that mapped population responses onto circular direction targets (cos *θ*, sin *θ*). Decoding performance was quantified by circular error, and decoder confidence was estimated from resultant vector length (Fig. 6A). Decoding error decreased systematically with stimulus coherence and confidence increased with directional signal strength (Fig. 6B-C), confirming that population activity carries reliability-dependent directional information.

**Figure 6:**
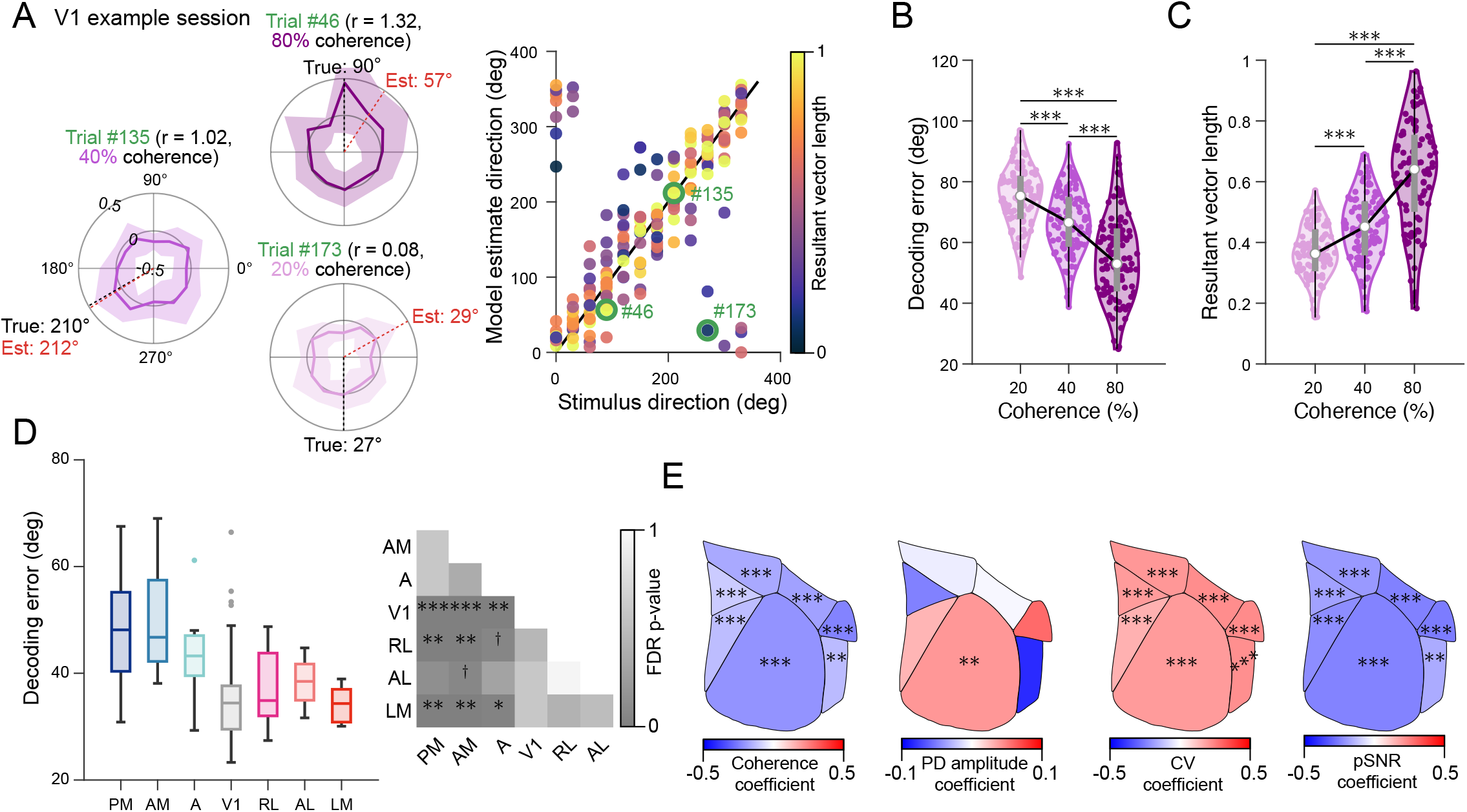
Single-trial population decoding errors scale with strength of reliability encoding features. **(A)** Left, performance of population-decoder on example single trials with different values of resultant vector length (r). Single-trial population activity, shown with shaded standard deviation of responses across neurons sharing a preferred direction. Black dashed line, true stimulus direction; red dashed line, decoder-estimated stimulus direction. Right, decoding performance for all trials within a single session (*n* = 180, non-zero coherence trials; V1 population), colored by resultant vector length. Black line, perfect decoding; example trials from left, green outlines. **(B)** Population decoding performance improves with stimulus coherence. **(C)** Population resultant vector length (a measure of confidence) increases with stimulus coherence. **(D)** Decoding performance comparison between visual areas. **(E)** Coefficients of area-specific multiple linear regression models predicting decoding error from reliability encoding features. Statistics from (B) and (D) based on pairwise comparisons using LMM contrasts with FDR-correction for multiple comparisons. ^†^*P <* 0.1, \**P* < 0.05, \*\**P* < 0.01, \*\*\**P* < 0.001.

To compare decoding across V1 and HVAs, we sub-sampled the most responsive neurons from each area to a fixed set size (*n* = 75 responsive neurons per area; Fig. S6A). Decoding was consistently more accurate in V1 and lateral HVAs (RL, AL, LM) than in medial visual areas (PM, AM, A; Fig. 6D), mirroring the stronger population drive and more consistent responses to coherent motion (higher pSNR) observed in these areas (Figs. 5B,E). This pattern was conserved even when neurons were selected randomly rather than by SNR rank (Fig. S6B-C), confirming that the lateral advantage in decoding reflects genuine differences in population encoding rather than neuron selection.

We next asked which features of the population tuning curves extracted above best predicted single-trial decoding error. We used area-specific multiple linear regression models with coherence and three population features as predictors: PD amplitude, circular variance, and pSNR. Opposite direction amplitude was omitted as a predictor given its close correspondence with PD amplitude across areas (Fig. 5B-C). As expected, coherence was a significant predictor of decoding error across all areas (Fig. 6E). However, on a trial-by-trial basis within a coherence level, pSNR and circular variance were stronger predictors of decoding accuracy than PD amplitude (Fig. 6E). This dissociation suggests functionally distinct roles among reliability-encoding mechanisms. Amplitude broadly tracks stimulus coherence, while the geometry of the population response, captured by pSNR and circular variance, more directly constrains the accuracy of single-trial directional estimates.

## DISCUSSION

Here, we show that the geometry of neural population activity encoding sensory reliability is conserved throughout mouse visual cortex. In both single neurons and populations, motion coherence modulates direction-tuned responses via coordinated changes in response amplitude, tuning width, and trial-to-trial variability. Although these features are differentially enriched across visual areas, they define a shared, multi-dimensional coding scheme for reliability. This organization argues against models in which reliability is represented by dedicated neuronal subpopulations or isolated features, and instead supports a unified code whose expression is locally scaled by circuit function and input drive. Consistent with this view, areas exhibiting stronger modulation along these dimensions support more accurate population-level decoding of motion direction, directly linking reliability-related structure to downstream perceptual readouts. Together, these findings constrain models of probabilistic inference by ruling out simplified or modular implementations of reliability signaling and instead favoring a multi-dimensional population-level geometry in which reliability emerges from coordinated changes in response structure. More broadly, this work establishes a foundation for future causal studies aimed at determining how reliability signals are selectively accessed to support perceptual decisions.

### Population decoding of motion is more accurate in lateral than medial areas

Converging anatomical and functional evidence supports the existence of two sub-networks specializing in the processing of object-related (ventral) and action-related (dorsal) vision (32, 31, 34, 53), although this organization is less segregated and more contentious in mice than in primates (32, 31, 34, 54). Within this framework, LM and AL are respectively considered the gateways from V1 to the ventral-and dorsal-like processing streams (55, 32, 53). However, we observed broadly similar motion direction and reliability encoding properties in AL and LM, at both the single-neuron and population levels (Figs. 4-6 and S5), suggesting the segregation of visual information along the two processing streams is not as well defined as in primates. Further, population decoding of motion direction was more accurate in lateral areas (AL, LM, RL) than in medial visual areas (PM, AM, A), despite the latter being frequently implicated in motion-related processing (31, 34, 32). Lateral areas also exhibited larger coherence-dependent increases in response amplitude from 0% to 80% coherence than medial areas (Fig. 2J), suggesting enhanced sensitivity to the specific motion features of the presented stimulus. Retinotopic alignment to the stimulus likely contributed to this pattern as well, since the centrally-presented stimuli preferentially engaged lateral areas through stronger retinotopic alignment. Together, these results suggest that the functional specialization across visual areas cannot be fully captured by a simple dorsal-ventral dichotomy in the mouse, but additionally depends on the interaction between stimulus statistics and receptive field alignment.

### Paradox of medial areas

If single neurons in medial visual areas are conventionally more selective for visual motion, why does their population code support worse decoding (Fig. S5; (32, 31))? The apparent paradox suggests a dissociation between single-neuron selectivity and population-level information. In medial areas, neurons showed narrower tuning curves and stronger suppression of opposite direction responses (Figs. S5B-C), features of increased directional selectivity. This pattern was also present at the population level, albeit marginally (Figs. 5C-D). However, medial areas simultaneously exhibited weaker population drive (lower preferred-direction response amplitudes; Fig. 5B) and higher variability (smaller pSNR; Fig. 5E). Fewer neurons were driven by the stimulus, and those that responded were less reliable. As a result, enhanced stimulus selectivity at the level of individual neurons was insufficient to drive improved population discriminability (56, 57). Population-level encoding of motion direction depends jointly on tuning sharpness, response amplitude, and variability. The advantages seen by medial areas in the former were largely outweighed by disadvantages in the latter two.

### Mechanistic origin of coordinated modulation

Across visual cortex, reliability-dependent modulations were strongly coupled. Neurons that exhibited coherence-dependent changes in tuning sharpness or response variability almost always showed concurrent changes in response gain, whereas isolated modulation of any single feature was comparatively rare (Figs. 3-4). Divisive normalization offers a strong candidate mechanism to explain this coordinated modulation. Broadly, divisive normalization is a canonical computation in visual cortex, originally characterized in the motion-sensitive visual circuits of primate MT (58, 59, 60). Under divisive normalization, a neuron’s response to a particular stimulus is scaled down (divided) by the summed activity of the surrounding population of neurons, with the relative contributions of neurons tuned to different stimulus features giving rise to inhibition from opposite direction (180° offset) stimulation (25, 61, 62). As motion coherence increases, the ratio of preferred to non-preferred direction drive increases, simultaneously amplifying responses in the preferred direction and suppressing responses elsewhere. Thus, divisive normalization naturally links changes in gain and tuning sharpening, a phenomenon that has been observed experimentally in primate MT (25, 37, 60) and is consistent with the opposite-direction suppression we observed in medial areas (Figs. 5C). Reduced trial-to-trial variability with increasing coherence is also consistent with normalization frameworks, in which stronger net input reduces the relative contribution of internal noise (41, 40). Importantly, divisive normalization is not mutually exclusive with known top-down contributions from higher areas (63, 64, 34, 65). Feedback from these areas could, in principle, sharpen reliability signals in early sensory cortex, though our passive viewing design does not allow us to disambiguate feedforward and feedback contributions. Distinguishing these signals, perhaps through targeted optogenetic inactivation of feedback pathways, represents an important target for future work.

## Acknowledgments

We thank the members of the Dadarlat and Sereno labs for helpful discussion and insightful comments on this manuscript. This work was supported by funds from the Indiana Clinical and Translational Sciences Institute (CTSI), funded in part by NIH Award Number UL1TR002529 (M.C.D., A.B.S), NSF HDR Grant 2117997 (M.C.D.), a Health of Forces grant from Purdue University (M.C.D, A.B.S.) and the National Defense Science and Engineering Graduate Fellowship (E.M.E.).

## Author contributions

E.M.E. - Conceptualization, Data Curation, Formal Analysis, Investigation, Methodology, Software, Visualization, Writing – original draft.

M.H.L. - Investigation, Writing - reviewing and editing.

A.B.S., M.C.D. - Conceptualization, Funding Acquisition, Methodology, Project Administration, Supervision, Writing - reviewing and editing.

## METHODS

### Resource availability

#### Lead contact

Further information and requests for resources and reagents should be directed to and will be fulfilled by the lead contact, Maria Dadarlat.

#### Materials availability

This study did not generate new unique reagents. The data and custom analysis code used here will be made publicly available upon acceptance of this manuscript.

### Experimental model and subject details

#### Animals

All animal procedures and experiments were approved by the Institutional Animal Care and Use Committee at Purdue University. A total of 9 GCaMP6s-expressing adult mice were used in this study. Eight mice contributed neural data (4M; Table S3) and 5 contributed behavioral data (2M; Table S4), with 4 mice contributing to both. GCaMP6s expression in putative excitatory neurons was induced by crossing of the following mouse lines: TRE-GCaMP6s (RRID:IMSR_JAX: 024742) x CaMKII-tTA (RRID:IMSR_JAX:007004). Mice were housed on a standard 12 h light/dark cycle with *ad libitum* access to food and water. Experiments were performed during the light cycle.

### Experimental procedures

#### Surgical procedures

All mice (*n* = 9) underwent two surgical procedures in preparation for experiments: (1) implantation of a titanium headplate for head-fixation and (2) implantation of a cranial window to provide optical access for 2-photon imaging. For all surgical procedures, mice were anesthetized using isoflurane in oxygen (3% induction, 1.5%-2% maintenance) and given a subcutaneous injection of meloxicam (5-10mg/kg) for postoperative analgesia. Body temperature was maintained using a homeothermic heat pad. Ophthalmic ointment was applied to provide eye lubrication. Depth of anesthesia was confirmed throughout surgeries by continued absence of a toe-pinch reflex.

For the headplate implantation, an additional subcutaneous injection of 0.03 mL lidocaine (0.5% solution) was delivered to the scalp to provide localized anesthesia. A small incision was then made and an area of skin excised over the skull. Remaining fascia was cleared and the skull surface cleaned and dried. To improve dental cement adhesion, the temporal muscles on both sides of the skull were retracted using forceps and the surface of the skull was etched. A thin layer of dental cement (C&B Metabond) was applied over the exposed surface of the skull and in the pockets created by retracted muscle. A circular titanium headplate (8mm inner diameter) was then attached over the posterior portion of the right cortical hemisphere using dental cement.

A minimum of 3 days later, mice underwent a second surgery to implant a 5mm cranial window over V1 to allow optical access to the brain. Prior to surgery, mice were given an additional subcutaneous dose of dexamethasone (2.0mg/kg) to reduce cerebral edema. A 5mm craniectomy was then made over V1 using a 0.5mm steel drill bit, as far lateral and caudal as was possible without compromising the integrity of the implant. The skull surface was periodically cooled by irrigating with chilled artificial cerebrospinal fluid (ACSF). A 5mm glass window was placed over the exposed brain and secured to the surrounding skull using dental acrylic (Lang Dental Ortho-Jet). Following each surgery, mice were then allowed to recover on a homeothermic heating pad and administered meloxicam (5-10mg/kg) every 24 hours for 2 days post-operation to reduce pain. Mice were given at least 3 days to recover from the cranial window implant before habituation to head-fixation.

#### Visual stimuli

All visual stimuli were generated using the PsychoPy® library in Python (66). For passive-viewing experiments, stimuli were presented for 2s, alternating with a gray-screen inter-stimulus interval lasting 4s. Stimuli were presented on a flat LCD monitor (38.6 × 61.2 cm, 1440 × 2560 pixels, 60 Hz refresh rate) positioned bilaterally 25cm in front of the eyes, such that it spanned approximately 68° in azimuth and 100° in elevation.

Random dot kinematograms (RDKs) consisted of white dots presented on a black background. Each dot had a diameter of 3.8°, moved across the screen with a speed of 25°/s, and had a lifetime of 10 frames (167ms; (32, 19, 31, 20)). The number of dots was set such that dots occupied 12.5% of the screen.

On each trial, a subset of dots–the *coherent* fraction (20, 40, 80%)–moved in one of 12 randomly assigned directions (0°/s - 330°/s in 30° increments), while all others moved randomly. Each coherence and direction combination was repeated five times, using an identical pattern of dot motion between repeats. Additionally, each session contained five trials in which there was 0% coherent motion, creating a null directional signal. In total, mice viewed 185 RDK trials per session, with order randomized between sessions. Each mouse participated in 2-6 sessions (Table S3).

During a subset of sessions, retinotopic maps were generated by sweeping a bar across the monitor that was spherically corrected to simulate spherical coordinates on the planar surface (33, 35, 31). The bar subtended 12° in the direction of motion and filled the monitor in the perpendicular direction. A contrast-reversing checkerboard pattern was presented within the bar to drive neural activity (0.04 cycles/° spatial frequency, 6Hz temporal frequency; (33)). Each trial consisted of a sweep of the bar in one of the four cardinal directions with a temporal period of 22.3s (33). Five trials were presented for each of the four directions with randomized order. Trials were separated by a gray screen period of 5s to allow stimulus-evoked activity to subside (48).

#### *In vivo* two-photon calcium imaging

Two-photon imaging was performed using a 2-photon mesoscope (Neurolabware). A mode-locked Ti:Sapphire laser (Coherent Chameleon Vision II) tuned to 920 nm was used to excite GCaMP6s expressed in layer 2/3 neurons. Emission light was collected by a Nikon 16X water-immersion objective (NA 0.8). All experiments were controlled using Scanbox software with a 512×796 pixel field of view sampled at 7.81 Hz. Imaging was performed at 1× magnification, such that each imaging window had an approximately 1.2 × 1.9mm field of view.

During imaging, mice were awake, head-fixed, and trained to sit in a custom 3D printed tube with a detachable top. Imaging was performed in mice trained to passively view visual stimuli. During imaging, we monitored the position and diameter of the left pupil and movement of the mouse at 60Hz frame rate with a video camera (DMK 37BUX287 cameras; Computar Telecentrix TEC M55 lens, pupillometry; TPL 0820 lens, behavior camera; (67, 68)). A high-powered infrared LED (Mightex SLS-0208-A) was used to provide additional illumination for imaging. To prevent direct contamination of the photomultiplier tube by light from the stimulus monitor, a custom light block was placed around the base of the objective.

#### Behavioral training

The five mice contributing behavioral data to this study all underwent the training protocol described below (Table S2). The remaining four mice used for imaging were trained on a related motion discrimination task and are not included in the behavioral analysis. Prior to behavioral training, all mice underwent the headplate implantation surgery described above. To motivate task engagement, mice were maintained on a citric acid water protocol (0.5-5% concentration), providing ad libitum fluid access, with daily weight monitoring to ensure animals remained above 80% of baseline body weight (69, 43). Training proceeded in three stages (43). Mice were first habituated to handling and head-fixation to the same 3D printed tube used in imaging (7 days). Then, mice were introduced to the task structure over three days, during which coherent motion stimuli at 100% coherence were presented in the binocular field of view, an auditory “go” cue was played, and visual flow was gradually slowed. Mice received a liquid reward (strawberry Nesquik) upon cessation of stimulus flow. Finally, mice were trained on the full two-alternative forced choice (2AFC) task, learning to rotate a wheel with their forepaws to report the horizontal motion direction of RDK stimuli at variable coherence levels. Mice initiated trials by holding the wheel steady for 300 - 700 ms. An RDK stimulus then appeared with global motion in the 0° (rightward) or 180° (leftward) direction. Behavior stimuli had the same parameters as used for imaging: each dot was of size 3.8° and moved with an initial speed of 25°/s (white dots, black background) (32, 20, 19). After a passive viewing period of 500-1000ms, a 100-ms tone signals trial onset. Wheel rotation opposite to the stimulus direction (e.g., counterclockwise rotation for rightward motion), slowed dot velocity while wheel rotation matching stimulus motion increased dot velocity. Mice were given a liquid reward for “zeroing” dot velocity. Incorrect responses (wheel movements doubling initial dot speed) or failure to respond within 10s triggered an aversive white noise stimulus. Correction trials were used to discourage directional bias. Mice were considered proficient at this task upon achieving ≥75% accuracy over three consecutive days of training, typically requiring approximately 20 training days.

#### Behavioral performance

Following training, mice underwent a minimum of 4-5 testing sessions at the full coherence range (0-100%) to establish a stable psychometric curve (19, 20, 70). During testing, stimuli were presented at seven coherence levels (0, 20, 40, 60, 80, 90, 100%) with motion direction randomized across trials. Trials were split into alternating easy (80, 90, 100% coherence) and hard (0, 20, 40, 60, 80% coherence) blocks to help maintain engagement throughout the session. Sessions were included in analysis if mice achieved ≥70% correct on high coherence trials (≥80% coherence), where timeout trials were treated as incorrect responses for this criterion. For the psychometric analysis, timeout trials were excluded from accuracy calculations, and choice accuracy was computed as the proportion of correct responses at each coherence level. A psychometric function was fit to the mean accuracy data (averaged across sessions and mice), using a cumulative Gaussian (19):

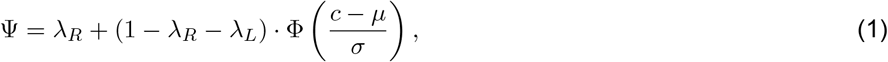

where *λ*_*R*_ and *λ*_*L*_ are leftward and rightward lapse rates, respectively, *µ* is the bias term, *σ* is the threshold parameter, and Φ is the standard normal cumulative density function (CDF). Parameters were estimated by minimizing the negative log-likelihood function using MATLAB *fminsearch*. The coherent motion perception threshold was defined as the coherence level yielding 70% correct performance (19, 20).

### Data analysis

#### Processing of calcium imaging data

Calcium imaging data was processed using the Suite2p toolbox (44), using default motion and neuropil correction settings, with an indicator decay timescale of 1.25 (45). We retained ROIs with cell probability *≥* 50% and fluorescence trace skew *≥* 0.5.

#### Stimulus responses

Unless otherwise stated, we defined the stimulus response for each neuron as the mean deconvolved fluorescent activity (Suite2p variable “spikes”) over the 16 frames (2s) following stimulus onset.

#### Retinotopic mapping

To generate retinotopic maps, we used neuropil responses to presentations of the sweeping bar stimulus described above (45). Raw neuropil fluorescence responses were averaged across stimuli of each direction. For each neuropil mask, we then computed the 1D Fourier transform of the four direction-averaged traces and extracted the phase of the first harmonic (71, 33). Maps were spatially smoothed by binning neuropil into 0.1 mm × 0.1 mm spatial grids based on the relative positions of their associated cell bodies within the imaging field-of-view (FOV). Retinotopic (phase) maps were phase-wrapped to ensure smooth transitions across the grid (31). To cancel the stimulus delay due to GCaMP6s rise time, we subtracted maps from stimuli moving in opposite directions (upward vs. downward, forward vs. backward) (71, 31). Finally, we remapped phase angles to visual degrees based on location of the stimulus screen relative to the mouse (Figure S1).

#### Registration to Allen Common Cortical Framework (CCF)

The purpose of registering retinotopic maps to the Allen CCF was to assign a standardized location to each neuron recorded from different imaging sessions and different mice. This procedure was modified from (46, 72).

For each mouse, we aligned the FOVs from each imaging session using a set of rigid transformations based on surface vasculature patterns and a high resolution image of the cranial window taken after surgery. This generated a location for each FOV in stereotaxic coordinates. We then applied this mapping to align the altitude and azimuth maps from each imaging session. In cases where imaging sessions overlapped, the median phase angle of neurons within overlapping grids was kept. Visual field sign was calculated by taking the sine of the angle between the gradients of the altitude and azimuth retinotopic maps (33). The map of field sign was then registered to the Allen CCF by aligning the border between V1, PM, and AM to the top-view CCF area contours (available from https://github.com/Zapit-Optostim/AllenAtlasTopDown/releases/tag/v0.1.4) based on established reference field sign maps (Fig. S1; (33, 46, 48)) to generate a stereotaxic coordinate-to-CCF transformation function for each mouse.

In cases where the field sign map was insufficiently resolved, the CCF alignment transformation was refined using the established reversal of azimuth phase along the lateral V1 border that was well-conserved in our dataset (33, 48).

To register each individual neuron to the CCF, we took the location of each neuron within the imaging FOV (center of mass from Suite2p; (44)), converted FOV-centric locations to stereotaxic coordinates, and transformed to the CCF using the mouse-specific stereotaxic coordinate-to-CCF transformation function (Figure S1).

#### Visual area parcellation

In our imaging experiments, we tiled our FOVs across posterior cortex to sample neurons across both primary and higher visual cortical areas (Figs. 2F,H-J, 4B). In certain analyses, neurons were grouped into seven distinct visual areas: V1, PM, AM, A, RL, AL, and LM (Figs. 4C-H, 5, 6). Area parcellations were achieved by separating neurons according to the CCF area contours using the MATLAB *polyshape* function (Table S1).

### Quantification and statistical analysis

Statistical analyses were performed in MATLAB R2023b (MathWorks).

#### Fitted tuning curves and SNR

To identify neurons with reliable visual responses, we used a previously established SNR method (Fig. S2; (45)). Tuning curves were estimated for each neuron across non-zero coherence levels using a Fourier basis expansion. Specifically, stimulus direction *θ* (radians) was represented using *n*_*basis*_ = 5 sine and cosine basis functions:

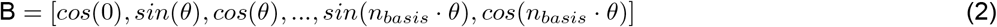

Linear regression was performed from the basis matrix *B* to the neural responses, yielding a fitted tuning function *f*_*n*_(*θ*). SNR was computed as ratio of the variance of tuning curve (*f*_*n*_(*θ*); signal) to the variance of the residuals (noise).

To compute coherence-specific *partial* SNR, the same procedure was used, but a separate tuning curve (*f*_*n,coh*_(*θ*)) was fit to responses on the set of trials with that coherence.

#### Selection of reliably responsive neurons

To identify neurons that reliably responded to the visual stimulus, we quantified stimulus-related response reliability using SNR (as described above) and assessed statistical significance relative to a neuron-specific scrambled null distribution.

For each neuron, the “true” SNR was computed on all non-zero trials as described above. To assess whether the observed SNR exceeded chance levels, we generated a neuron-specific null distribution by randomly permuting each neuron’s trial-by-time response matrix (100 permutations). This procedure preserved the original distribution of activity values while destroying the alignment between neural responses and stimulus identity. For each permutation, the mean response was recomputed for each trial and used to calculate a scrambled SNR using the same procedure as for the true data (45).

Neurons were classified as “reliably-responsive” to the visual stimulus if their true SNR exceeded the 90th percentile of the scrambled SNR null distribution. Key results were unchanged when more stringent thresholds (e.g., 95th percentile) were applied, although the number of neurons retained for analysis was reduced. We therefore report results using the 90th percentile threshold to retain sufficient statistically power given that comparison to single-neuron null distributions is already conservative. This percentile-based thresholding provided a non-parametric, neuron-specific criterion for significance that did not require assumptions regarding the shape of the null SNR distribution.

All SNR values, scrambled distributions, significance thresholds, and binary responsiveness labels were computed separately for each session and mouse. For population-level analyses, SNR values and responsiveness labels were pooled across mice and sessions.

#### Circular variance

To measure the degree of selectivity of each neuron to RDK motion direction, we calculated the circular variance of the response. Circular variance (CV) allows for a fit-less quantification of tuning selectivity (50).

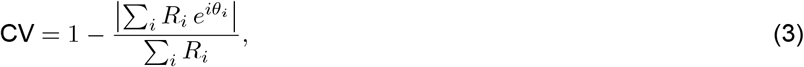

where *R*_*i*_ is the neural response at each stimulus direction *θ*_*i*_ (radians). By this definition, an exceptionally directionally-selective neuron that responds exclusively to a single direction, will have a circular variance of zero, whereas a cell lacking directional tuning that responds equally to all directions has a circular variance of one. Circular variance was computed using *Circular Statistics Toolbox* available from Mathworks.

#### Amplitude modulation index (AMI)

To determine the change in response amplitude between high (80%) and low (20%) coherence, we computed a normalized modulation index (AMI) as:

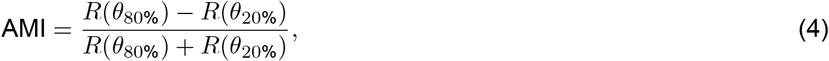

where *R*(*θ*_80%_) is the trial-averaged neural response to stimulus direction *θ* at 80% coherence. AMI was separately calculated for preferred direction (PD) and opposite direction (OD) responses. Amplitude modulation was quantified using a modulation index to normalize for differences in absolute response magnitude across neurons.

#### Selection of significantly modulated neurons

We computed three reliability encoding mechanisms: modulation of response amplitude (PD and OD), modulation of circular variance, and modulation of SNR. For each neuron, we quantified the amplitude modulation as described above (see Amplitude modulation index; AMI). Because circular variance and SNR are zero-bounded measures, modulation was defined as the raw change in CV or SNR from high-coherence (80%) to low-coherence and (20%).

We then performed a neuron-specific shuffling analysis to determine whether the observed changes exceeded those expected from sampling variability alone. For each neuron, trials from non-zero coherence conditions were pooled, preserving the joint distribution of response amplitudes and stimulus directions while removing the original association between coherence level and response structure. Trials were then randomly assigned to either the high-or low-coherence condition.

Features (response amplitude, CV, and SNR) were then recomputed for each shuffled group. The shuffled modulation was calculated as the AMI or raw change between the shuffled group features. Shuffling was repeated 100 times per neuron to generate a null-distribution of changes expected under the hypothesis that changes arise from sampling noise alone.

For each neuron, the “true” value of each modulation was compared against its shuffled neuron distribution. Neurons were classified as exhibiting a significant change if the absolute value of the true change exceeded the 90th percentile of the shuffle distribution.

Shuffle distributions, true changes, and significance labels were computed separately for each session and mouse. For visualization and summary statistics, results were pooled across mice and sessions. Population summaries were restricted to neurons previously classified as stimulus-responsive based on the SNR criteria (see *Selection of reliably responsive neurons*).

#### Population tuning curves

To generate population tuning curves, responsive neurons within each brain area recorded during a single imaging session were sorted according to their preferred directions as identified by the fitted tuning functions. On each trial, responses of neurons sharing a preferred direction were averaged to generate single-trial population tuning curves (1, 7, 73, 5).

Accordingly, 185 population tuning curves were generated for each visual area in which neurons were recorded during an imaging session (Fig. S4A). However, depending on the number of neurons recorded from that visual area and the distribution of preferred directions of recorded neurons, not all population tuning curves were complete. Incomplete tuning curves were considered in evaluations of preferred and opposite direction population amplitudes, but only tuning curves which contained a value for all 12 directions were used in calculating population CV and SNR.

#### Population SNR (pSNR)

The SNR method used at the single-neuron level measures the consistency of a neuron’s response across repeated trials of the same stimulus (45). To extend this concept to the population level, we developed single-trial, bounded population SNR metric, termed pSNR, that quantifies how closely the population response on a given trial matches the expected response for that stimulus condition.

Intuitively, if the population reliably encodes stimulus direction, repeated presentations of the same stimulus should evoke similar population tuning profiles across trials. Deviations from this expected response reflect trial-to-trial variability in the population code.

To quantify this consistency, we first circularly shifted each population tuning curve so that responses were aligned to a common stimulus direction of 0°. For each coherence, we then generated a leave-one-out population template by averaging the responses from all other trials.

For a given trial, we found the optimal scalar projection (g) of that trial response onto the population-average template:

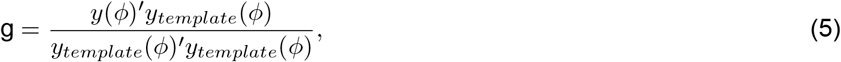

where *y*(*ϕ*) is the population response on a particular trial as a function of preferred direction (*ϕ*), *y*_*template*_(*ϕ*) is the leave-one-out population template associated with that trial.

We then obtained the predicted trial response (*ŷ* (*ϕ*)) by multiplying the population template, *y*_*template*_(*ϕ*) by the projection factor, g, and calculated the residuals (*r*(*ϕ*)) between the true and predicted responses for that trial. Population SNR was then calculated as the ratio of the square norm of the predicted response to the squared norm of the residual:

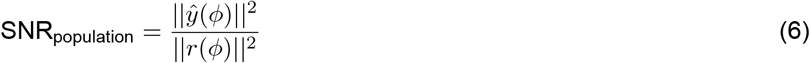

To obtain a bounded metric between 0 and 1, we calculate pSNR as:

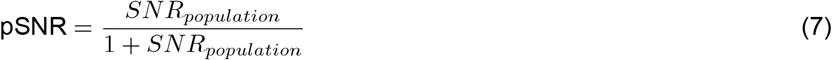

This metric quantifies the fraction of the population response energy that lies parallel to the population template, providing a measure of how strongly the population response on a particular trial aligns with the canonical response profile for that stimulus condition.

#### Population tuning modulation

Because population tuning curves were computed from single trials, we could not directly compute modulation as changes from average 80% coherence to average 20% coherence values as done for single neurons. Instead, modulation was computed for each mechanism (gain, tuning width, pSNR) by subtracting the trial-averaged responses at 20% coherence from each 80% coherence single-trial responses. As such, we obtained a modulation estimate for all 60 of the 80% coherence trials for each population. Note that for population tuning curves, gain modulation was directly estimated from this difference rather than a modulation index as population tuning curves were built from the z-scored deconvolved activity of single neurons, which is already a normalized measure.

#### Population direction decoder

For all single-neuron analyses, we took the coherence of the RDK to be a proxy for the reliability of the sensory signal, and used a correlational approach to evaluate how features of single-neuron responses co-varied with reliability. At the population level, we have access to single-trial response features of the population, which allows for estimation of the true trial-to-trial signal reliability. To measure single-trial information encoding, we trained a ridge-regularized (74), leave-one-out multivariate linear regression model to predict *Y* = [*cos*(*θ*), *sin*(*θ*)] where *θ* is the stimulus direction on a particular trial. The ridge-regularization parameter (*λ*) was tuned once per population using 5-fold cross-validation such that the each implementation of the leave-one-out decoder for the same population used the same optimal value of *λ*. For each trial, we were then able to calculate decoding error as the difference between the true stimulus value and the predicted stimulus value. Decoding error served as an estimate of the instantaneous reliability of the directional signal in the population.

Because the linear regression sees the response of each neuron in the population, to fairly compare between visual areas, it was necessary to control for the size of the population used in decoding. Progressively larger sample sizes were generated by adding neurons in reserve order of their SNR to ensure that areas were given the “best chance” at decoding and creating diminishing return on tuning quality with increasing population sizes. In all areas, decoding errors decreased with increasing population size (Fig. S6A). We therefore repeated this regression using a variety of population sizes and calculated for each area the point of diminishing return. We defined the median of these points across areas as the decoding population size (*n* = 75). Therefore, only imaging sessions in which at least 75 responsive neurons were recorded from a particular area were retained in decoding subsequent analyses.

#### Resultant vector length

To further support the decoding results, we additionally computed the Euclidean norm of the 2D predicted output vector *Y* = [*cos*(*θ*), *sin*(*θ*)]. This calculation returns the vector length (magnitude) which acts as a proxy for decoder confidence. If the population activity strongly supports a constant direction, the vector length will be close to one, indicating greater numerical stability of the angle estimate. Population activity that is noisy or distributed will shrink the vector towards the origin, leading to a vector length closer to 0.

#### Predicting decoding error using multiple linear regression

To quantify how population-level tuning properties relate to trial-by-trial decoding performance, we fit linear models to predict population decoding error from neural features. Analyses were performed separately for each visual area.

The dependent variable for the model was the population decoding error for each trial, measured in degrees. Predictor variables were population preferred direction (PD) gain, population circular variance, population SNR (pSNR), and stimulus coherence. Coherence values (20, 40, 80%) were treated as a continuous variable because this provided comparable explanatory power to categorical encoding (similar *R*^2^), while allowing a parsimonious model.

Before fitting the model, all predictors and the dependent variable were z-scored within each visual area to put coefficients on a common scale and enable comparison of effect sizes across predictors. We then fit a multiple linear regression of the form:

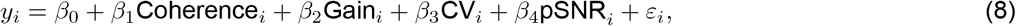

where *y*_*i*_ is the z-scored decoding error for population *i*; *β*_0_ is the intercept; *β*_1_–*β*_4_ are regression coefficients for each predictor; and *ε*_*i*_ *∼* 𝒩 (0, *σ*^2^) is the residual error. Regression coefficients and associated significance (FDR-corrected *P* values) were extracted for each predictor and visual area.

Across visual areas, the intra-class correlation coefficient (ICC) was low (mean ICC = 0.013, range = 0–0.02), indicating that mouse identity accounted for a small fraction of variance in decoding error. Consistent with this, results were qualitatively similar when including mouse as a random effect in the linear models. For simplicity, we therefore report results from linear regression models without random effects.

#### Statistical methods and models

The values of sample sizes and what they represent for each analysis can be found in the figure legends and related method sections. Reported *P* values are two-sided. Unless otherwise noted, the Benjamini-Hochberg procedure was used when multiple statistical tests were conducted simultaneously, such as comparing between visual areas, to control the false discovery rate (FDR) at 0.05 (MATLAB *mafdr*; (75)).

#### Generalized linear mixed effects models: Mechanism expression probability

This analysis is related to Figs. 4 and S3A. To test whether the probability of expressing each reliability encoding mechanism differed across visual cortical areas, we used generalized linear mixed effects models (GLMMs, MATLAB *fitglme*) with a binomial distribution and logit link function to model the binary neuron-level outcomes (mechanism expressed vs. not expressed).

These hierarchical models account for the nested structure of the dataset, in which neurons are sampled within imaging sessions, and sessions are nested within individual mice, precluding their treatment as independent observations.

For each mechanism, we fit a binomial GLMM with cortical area as a fixed effect and random intercepts for mouse identity and session nested within mouse identity:

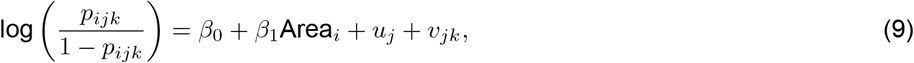

where *p*_*ijk*_ is the probability that a neuron in area *i*, mouse *j*, session *k* expresses the reliability encoding mechanism; *β*_0_ is the intercept on the log-odds scale; *β*_1_ is the fixed effect of cortical area; 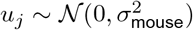 is a random intercept for mouse 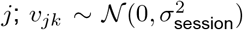 is a random intercept for session *k* nested within mouse *j*; and the binary outcome (expressed/not expressed) *y*_*ijk*_ *∼* Bernoulli(*p*_*ijk*_).

Pairwise contrasts between visual areas were evaluated by applying contrast vectors to the fixed-effect coefficient estimates. Effect sizes are reported as odds ratios (ORs) with 95% confidence intervals derived from the contrast estimates and their standard errors (Fig. S3).

To test whether mechanism identity predicted expression probability independent of cortical area, we additionally fit a pooled model including mechanism type as a fixed effect while keeping the previous fixed and random intercepts.

#### Linear mixed effects models: Between-area comparisons

These analyses are related to Figs. 5, S4, and S5. For continuous measures, we used an analogous linear mixed-effects framework (LMM; MATLAB *fitlme*) applied to the features extracted from tuning curves at 80% coherence and to the modulation of extracted features with coherence:

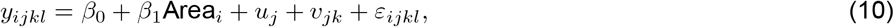

where *y*_*ijkl*_ is the *l*-th feature from brain area *i*, mouse *j*, session *k*; *ε*_*ijk*_*∼𝒩* (0, *σ*^2^) is the residual error; and all other terms are as defined above for the GLMM.

For single neuron area comparison analyses (Fig. S5), pairwise comparisons between areas were performed using linear contrasts on the fixed-effects coefficients and the resulting FDR-corrected *P* values were directly retained.

For area comparison analyses using population tuning curve features (Figs. 5 and S4), residuals were found to violate the heteroskedasticity assumption, so statistical conclusions were based on a cluster-level bootstrap resampling procedure (*n* = 1000 iterations) in which mice were resampled with replacement, session identifiers were reassigned to preserve nested data structure, and the LMM model was refit on the resampled dataset. For each pairwise area comparison, a contrast vector was applied to the full coefficient vector of each bootstrap model, generating an empirical null distribution of contrast estimates. Two-tailed bootstrap *P* values were computed by centering this distribution at zero and determining the proportion of resample contrasts whose absolute value exceed the observed contrast estimate.

#### Linear mixed effects models: Within-area comparisons

This analysis is related to Figs. 5F-I and S4F-I. An additional LMM was used to test for significant changes in population encoding features from 80% to 20% coherence (Figs. 5F-I and S4F-I). The same LMM structure was used, but statistical conclusions were based on a cluster-level bootstrap resampling procedure (*n* = 5000 iterations) comparing the model coefficient to zero.

#### Linear mixed effects models: Direction decoding performance

These analyses are related to Figs. 6 and S6. Two different LMM structures were used for statistical analyses of the results of population decoding. First, to evaluate trial-by-trial changes in decoding error and resultant vector length across coherence levels (Figs. 6B-C), an LMM was generated as in Eq. 10 with mouse and session nested within mouse as random effects, but the fixed effect of “Area” was replaced with “Coherence”. Thus, *y*_*ijkl*_ was the *l*-th feature from coherence *i*, mouse *j*, session *k*.

To compare decoding error across visual areas (Figs. 6D and S6), we performed analyses at the level of experimental sessions, treating each session as an independent sample of population activity within a given area. For each session, decoding error was averaged across trials to obtain a single estimate of population performance, reflecting that trials constitute repeated measurements of the same underlying population within a session and do not provide independent evidence about differences between areas. We therefore modified the LMM used in trial-level analyses (Eq. 10) to remove *v*_*jk*_, the random effect of session, because the present analysis operates on session-averaged data rather than individual trials, while retaining mouse as a random effect to account for repeated measurements across animals.

For all decoding analyses pairwise comparisons between coherences and areas were performed using linear contrasts on the fixed-effects coefficients and the resulting FDR-corrected *P* values were directly retained.

#### Linear mixed effects models: Between-coherence comparisons

A final LMM was used to test for changes in decoding error and resultant vector length across coherence levels (Figs. 6B-C). The same LMM structure with mouse and session nested within mouse random effects was used, but coherence was added as the fixed effect. Pairwise comparisons between coherence levels were evaluated using contrast vectors applied directly to the fitted model coefficients.

