## Supplemental Information for "A shared multi-feature population code for sensory reliability across mouse visual cortex"

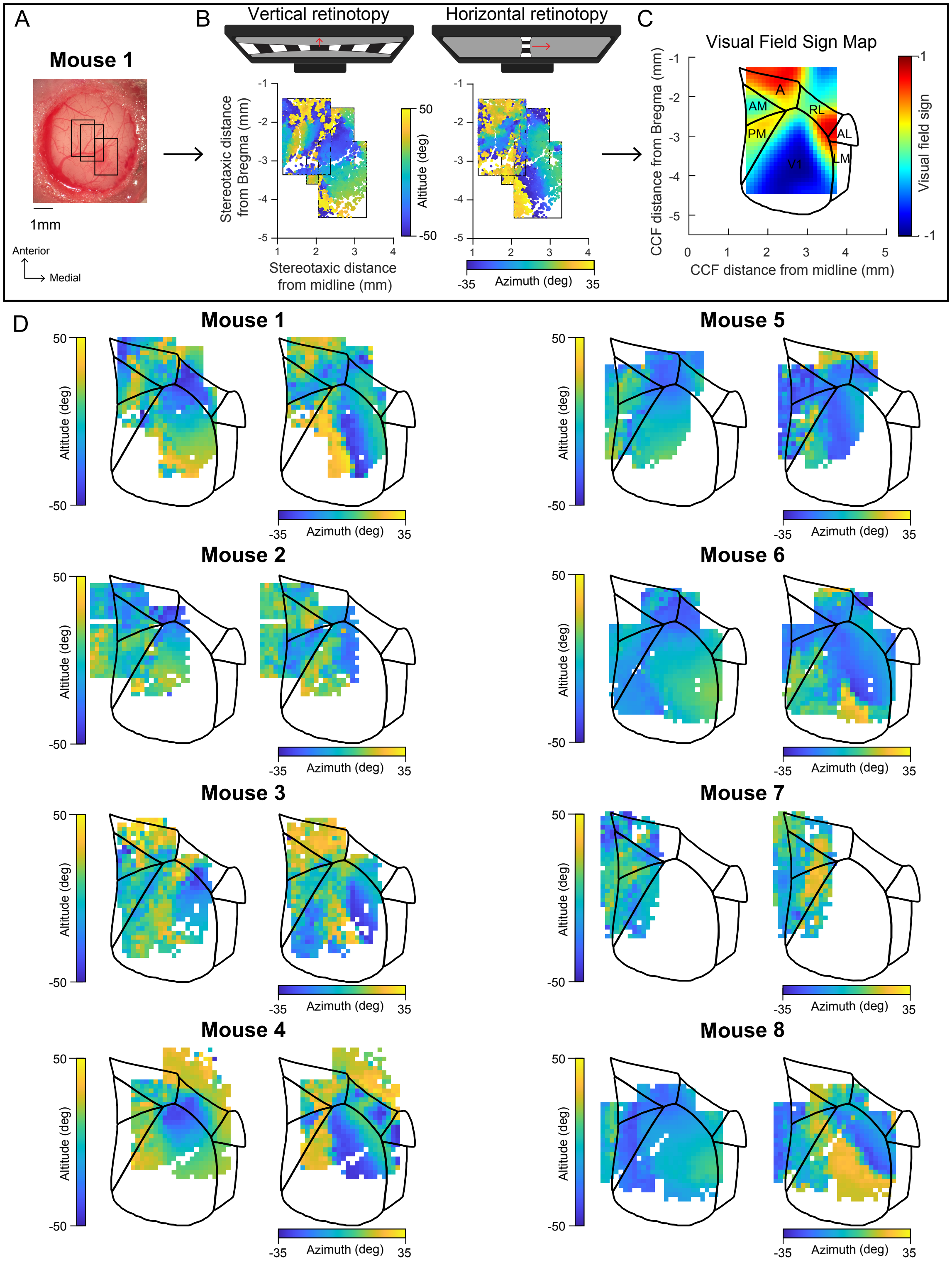


**Figure S1: Generation of retinotopic maps and registration of neurons to a common coordinate system; related to Figure 2. (A)** Overview image of the vasculature pattern within the cranial window for an example mouse. Boxes indicate the locations of all fields of view (FOVs) used for retinotopic mapping. Scale bar = 1mm. **(B)** Neuropil responses to the retinotopic mapping stimulus for the three FOVs from the mouse in (A), plotted in stereotaxic units. Top, screens depict retinotopic mapping stimuli, with a horizontally-aligned drifting bar moving vertically over the visual field (altitude mapping; left) and a vertically-aligned drifting bar moving horizontally across the visual field (azimuth mapping; right). Bottom, retinotopic maps showing preferred location of neuropil for altitude (left) and azimuth (right). Color bars indicate degree offset from center of visual field. **(C)** Registered visual field sign map (red, positive sign; blue, negative sign) for example mouse in (A). Visual field signs computed from retinotopic maps in (B). Visual area contours from the Allen Institute Mouse Common Cortical Framework (CCF). **(D)** Registered retinotopic maps for all mice used in passive viewing experiments (*n* = 8). Neuropil responses smoothed by binning to a 0.1 x 0.1mm spatial grid. Colors indicate preferred location of each pixel for altitude (left) and azimuth (right). Overlaid with Allen CCF area contours. Mouse 1 is example mouse from (A-C).

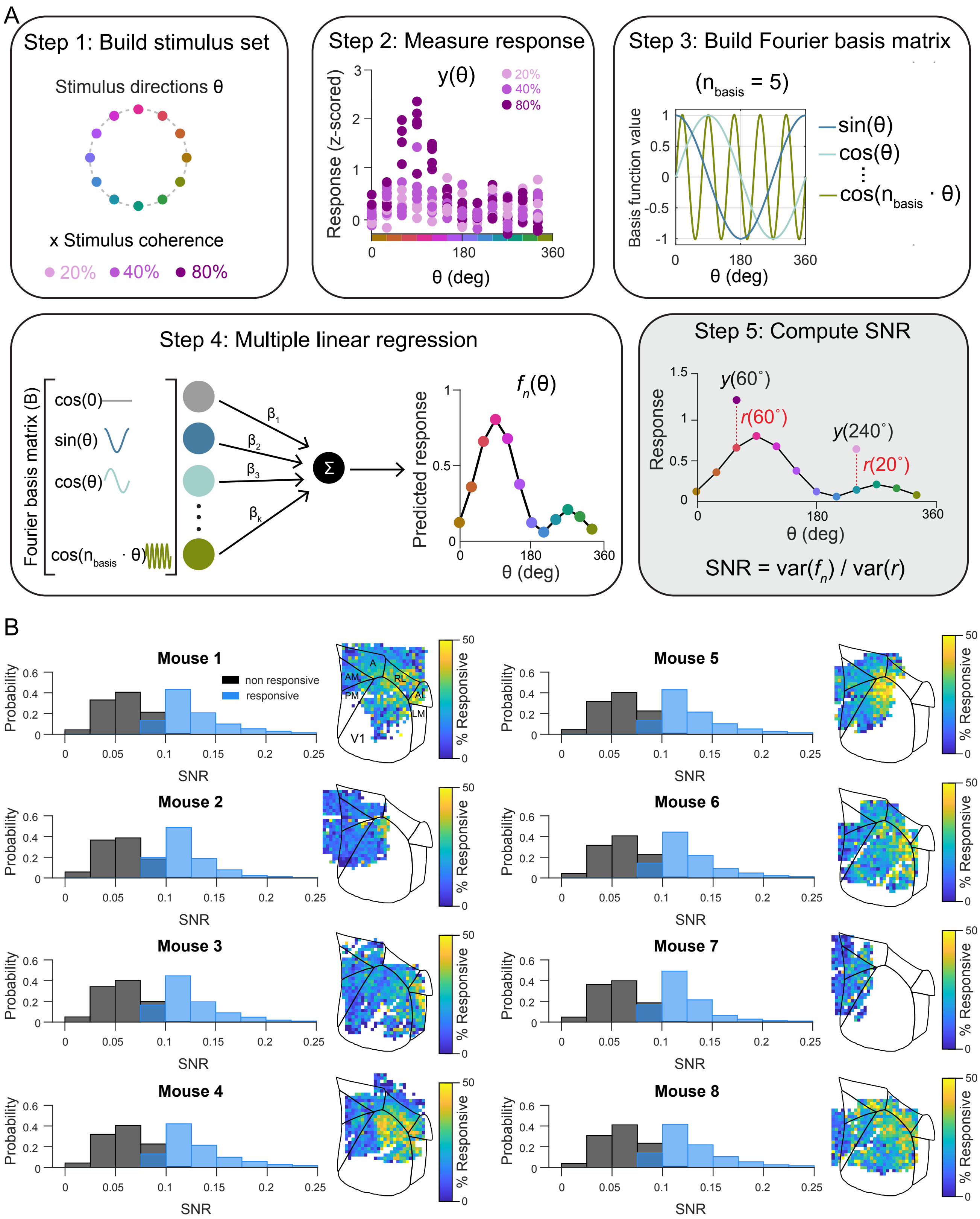


**Figure S2: Cross-animal consistency of responsive neurons using signal-to-noise ratio (SNR) metric; related to Figure 2. (A)** Framework for calculating SNR from neural responses to repeated stimulus presentation. SNR is calculated as the ratio of fitted tuning curve variance (signal; 5 sine and cosine basis functions) to residual variance (noise). **(B)** SNR distributions for each mouse. Responsive neurons (blue) are identified by determined by comparison to a per-neuron null distribution; non-responsive neurons are shown in gray (SNR below threshold). Right, spatial distribution of responsive neurons overlaid on the Allen Institute Mouse Common Cortical Framework (CCF) area map. Grid size: 0.1mm x 0.1mm.

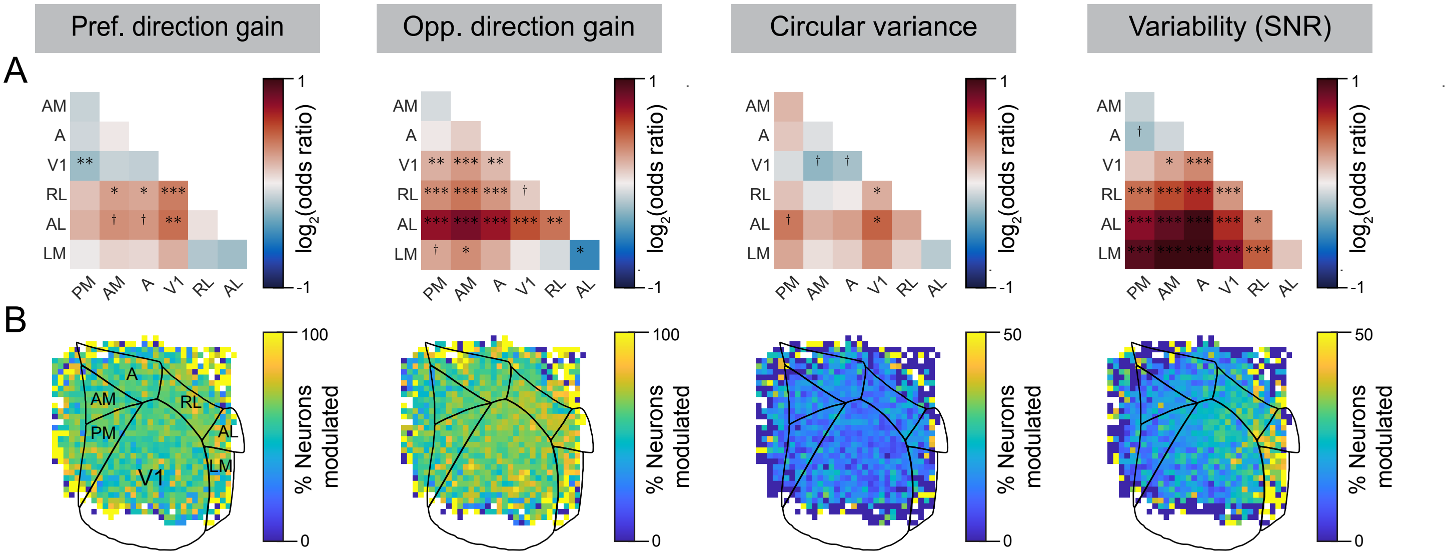


**Figure S3: Pairwise comparisons in the frequency of modulation mechanisms across visual areas; related to Figure 4. (A)** Odds ratios (ORs) for pairwise comparisons in encoding mechanism frequency (columns) using a generalized linear mixed effects model with a logit link function. ORs were log-transformed for visualization to produce a symmetric scale centered at zero. Symbols indicate FDR-correct $P$ values: $^{\dagger}P < 0.1$, *$P$ < 0.05, **$P$ < 0.01, ***$P$ < 0.001. **(B)** Spatial distribution of probability of mechanism use, expressed as fraction of responsive neurons implementing each encoding mechanism.

A

| Visual area | PM | AM | A | V1 | RL | AL | LM |
|---|---|---|---|---|---|---|---|
| # single-trial population tuning curves | 3780 | 2340 | 2880 | 6480 | 3240 | 720 | 540 |

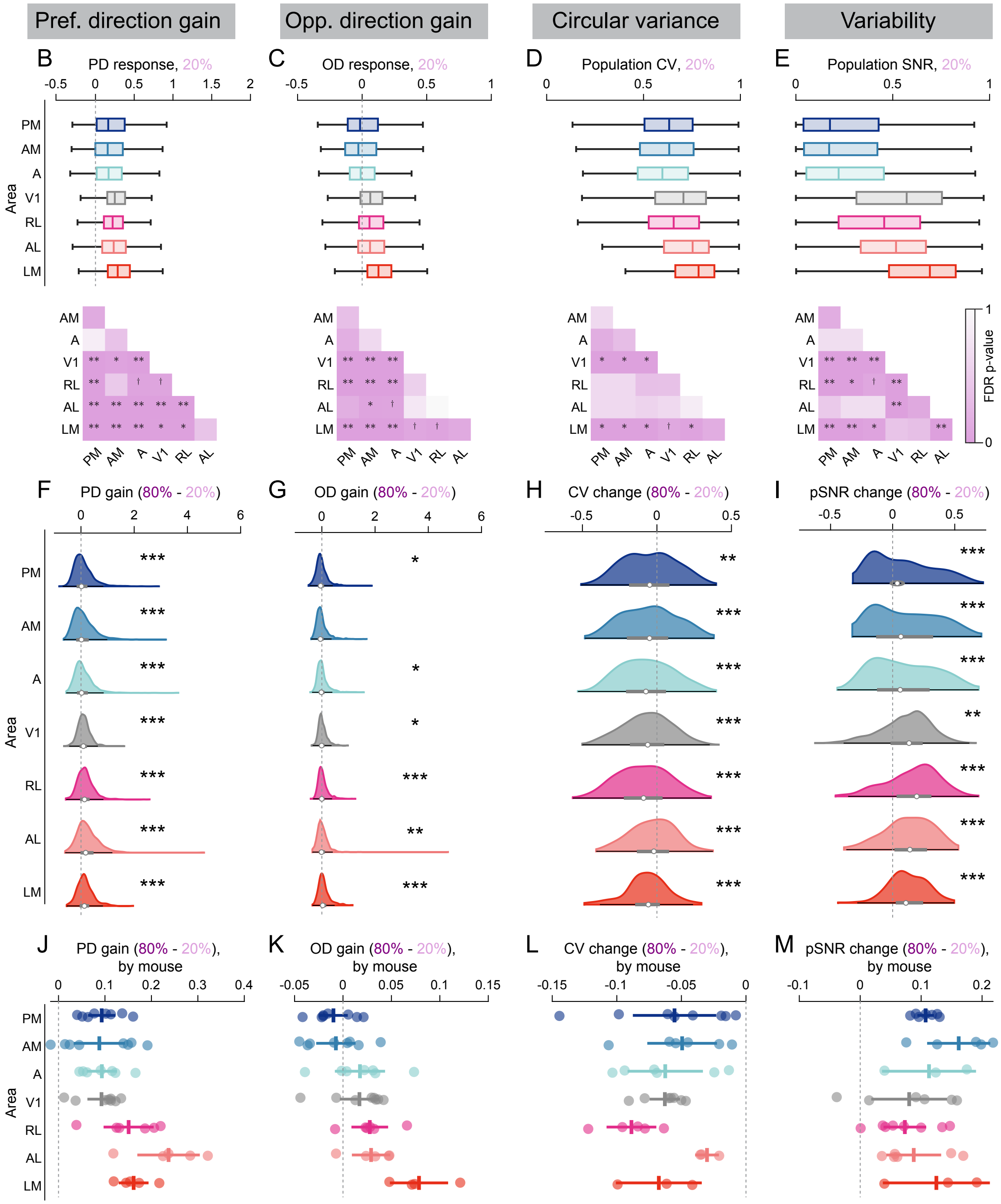


**Figure S4: Extended analysis of encoding mechanisms by neural populations; related to Figure 5. (A)** Number of single-trial population tuning curves generated per visual area ($n$ = 8 mice). **(B)** Preferred-direction (PD) population tuning curve response amplitude at 80% coherence. Bottom, pairwise comparisons between visual areas. **(C-E)** Same as (B) for opposite direction (OD) response amplitude (C), circular variance (D), and population bounded SNR (pSNR; E). **(F)** Full range of distribution of change in PD amplitude from 80% to 20% coherence. Circles, medians; gray bars, inter-quartile ranges. Dashed vertical line; zero change. Statistics indicate comparison of distribution to zero for each area. **(G-I)** Same as (F) for OD amplitude (G), circular variance (H), and pSNR (I). Bottom, pairwise comparisons of modulation across visual areas. **(J)** Mean change in preferred direction gain from 80% to 20% for each mouse, across all sessions. Colored lines, indicate group mean and 95% confidence interval; dashed gray lines, indicate zero change position. **(K-M)** Same as (J) for opposite direction gain (K), circular variance (L), and pSNR (M). All statistics indicate bootstrap LMM contrasts with FDR correction. $^{\dagger}P < 0.1$, $^{*}P$ < 0.05, $^{**}P$ < 0.01, $^{***}P$ < 0.001.

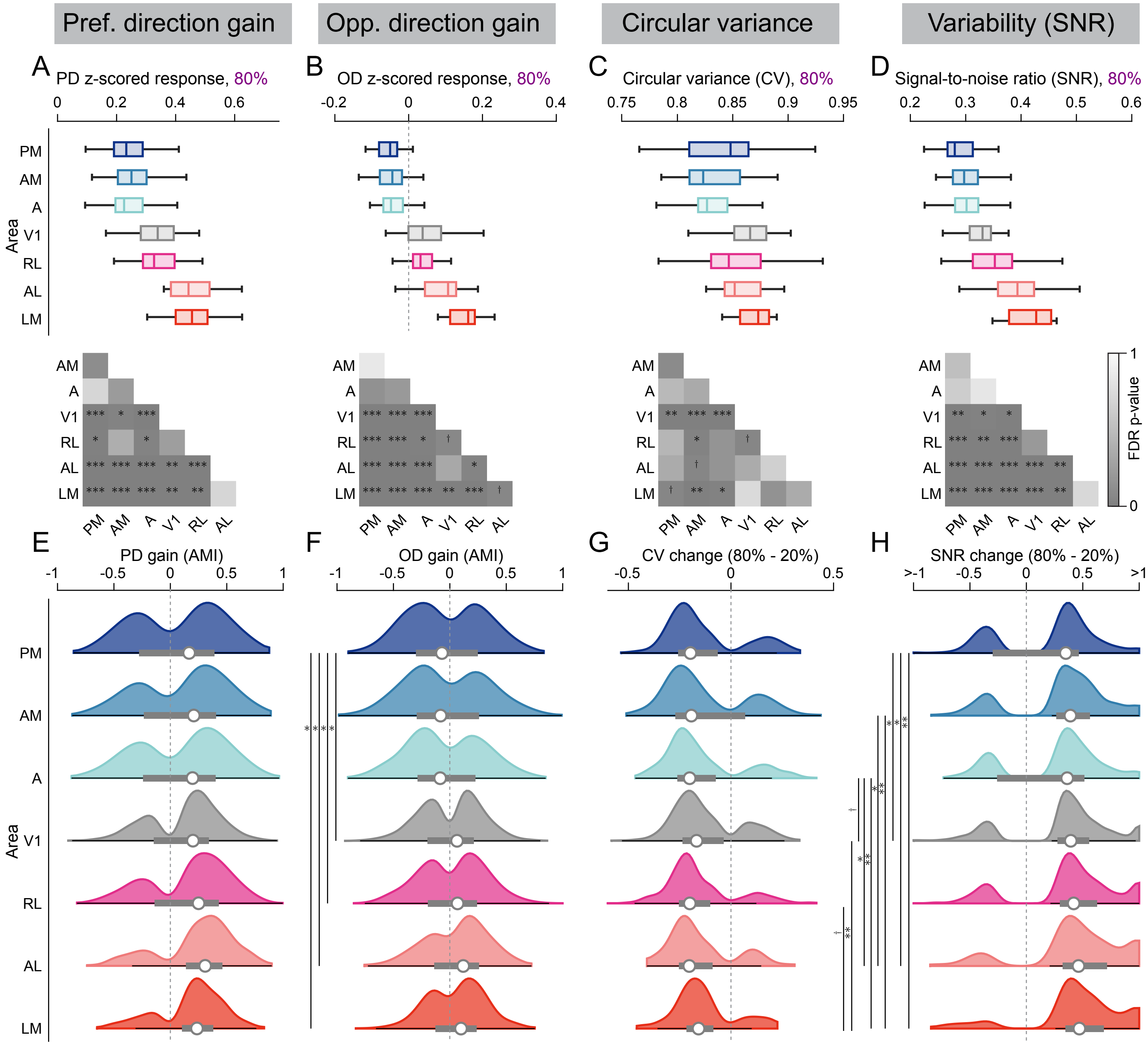


**Figure S5: Implementation of reliability encoding mechanisms by single neurons across visual areas; related to Figure 5. (A)** Preferred-direction (PD) amplitude of trial-averaged tuning curves for all responsive neurons at 80% coherence. Bottom, pairwise comparisons of amplitude at 80% coherence between visual areas. **(B-D)** Same as (B) for opposite direction (OD) amplitude (C), circular variance (D), and signal to noise ratio (SNR; E). **(E)** Distribution of PD gain among neurons identified as significantly-gain modulated (see **Figure 3**), expressed as the amplitude modulation index (AMI) between 80% and 20% coherence. Circles indicate distribution median; gray bars indicate inter-quartile range; black lines indicate whiskers. Dashed vertical line, zero change. **(F-H)** Same as (E) for OD AMI (F), change in circular variance from 80% and 20% (G), and change in SNR from 80% and 20% (I). Statistics reflect FDR-corrected $P$ values for pairwise comparisons between visual areas. $^{\dagger}P < 0.1$, $^{*}P$ < 0.05, $^{**}P$ < 0.01, $^{***}P$ < 0.001

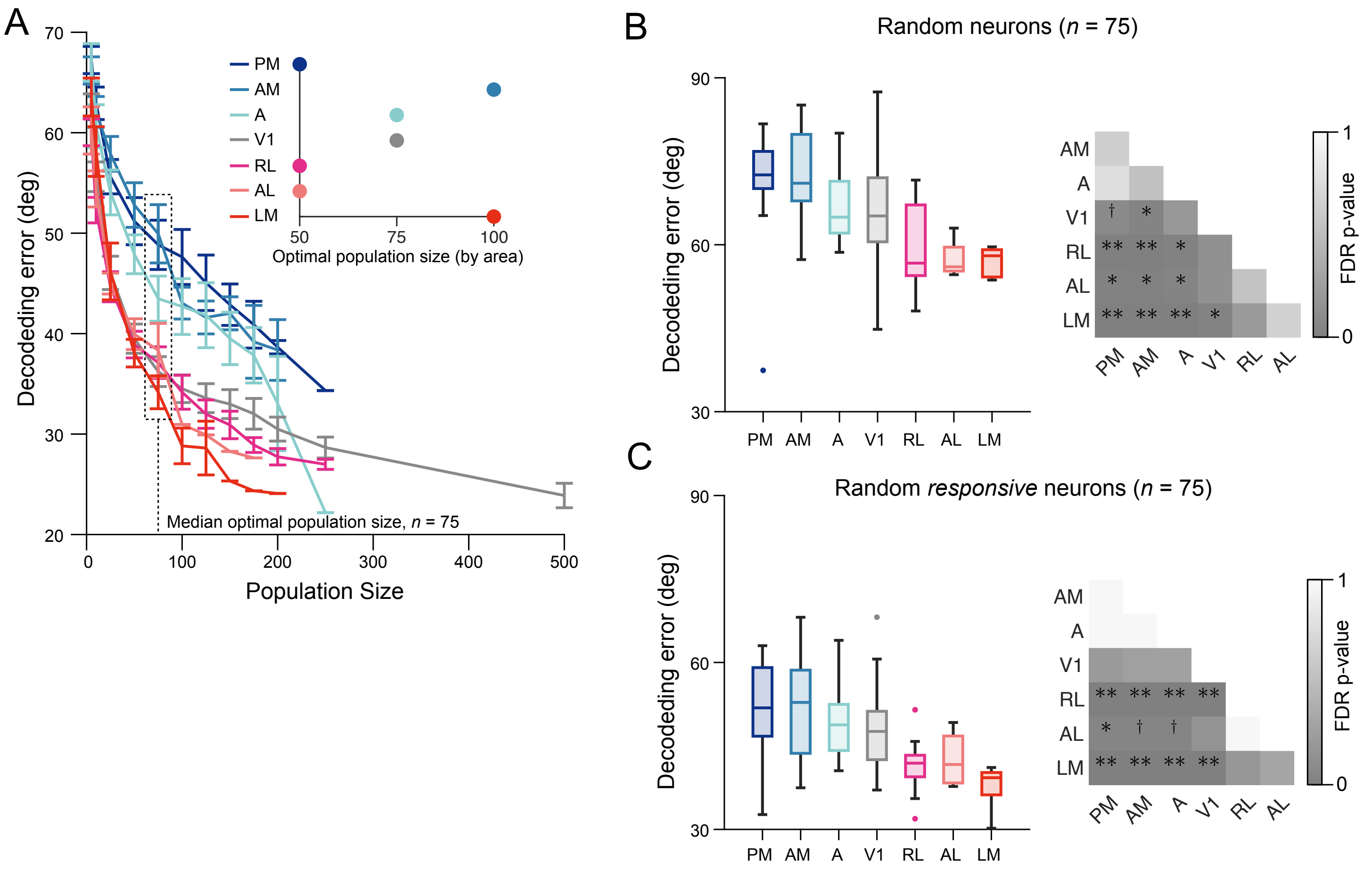


**Figure S6: Effects of sub-sampled population size on directional decoding; related to Figure 6. (A)** Decoding error as a function of population size for each visual area. Points indicate mean ± SEM across sessions achieving that population size threshold. Neurons were added in reverse SNR rank order, retaining the most responsive neurons in smaller populations. Inset, optimal population size per area, defined as the point at which decoding error improved by less than 0.1° per additional neuron. The median optimal population size across areas (dashed box), was used for decoding analyses. **(B)** Decoding performance using 75 randomly selected neurons per area without SNR restrictions. **(C)** Decoding performance using 75 randomly selected responsive neurons per area, relaxing the SNR rank constraint. For (B) and (C), statistics indicate FDR-corrected $P$ values for pairwise areas comparisons using LMM contrasts. $^{\dagger}P < 0.1$, *$P$ < 0.05, **$P$ < 0.01, ***$P$ < 0.001.

**Table S1: Summary of imaging sessions; related to Figure 2.** Neuron counts reflect recordings retained after skewness thresholding; responsive fractions reflect SNR-based reliability classification. Only areas with recorded neurons are reported per session. V1 = primary visual cortex; PM = posteromedial; AM = anteromedial; A = anterior; RL = rostrolateral; AL = anterolateral; LM = lateromedial.

| Mouse | Session | # Neurons (% responsive) | # Responsive neurons by area (% responsive) |
|---|---|---|---|
| 1 | 1 | 2955 (0.21) | V1: 1104 (0.24), PM: 197 (0.21), AM: 221 (0.24), A: 959 (0.18), RL: 368 (0.25) |
| 1 | 2 | 2775 (0.21) | V1: 2659 (0.20), RL: 111 (0.26) |
| 1 | 3 | 3813 (0.19) | V1: 1008 (0.22), PM: 227 (0.18), AM: 311 (0.21), A: 1472 (0.15), RL: 600 (0.23) |
| 1 | 4 | 3124 (0.24) | V1: 474 (0.27), A: 23 (0.22), RL: 1279 (0.29), AL: 183 (0.31), LM: 46 (0.37) |
| 1 | 5 | 3673 (0.18) | V1: 217 (0.27), PM: 554 (0.14), AM: 1293 (0.18), A: 1149 (0.19) |
| 1 | 6 | 3753 (0.28) | V1: 912 (0.37), PM: 251 (0.39), AM: 281 (0.31), A: 1512 (0.23), RL: 653 (0.25) |
| 2 | 1 | 2708 (0.19) | V1: 1283 (0.20), PM: 720 (0.18), AM: 414 (0.20), A: 252 (0.11), RL: 1 (0.00) |
| 2 | 2 | 2468 (0.14) | PM: 177 (0.19), AM: 238 (0.13), A: 84 (0.10) |
| 2 | 3 | 2925 (0.13) | V1: 1327 (0.14), PM: 762 (0.13), AM: 369 (0.09), A: 435 (0.13) |
| 2 | 4 | 3601 (0.11) | V1: 262 (0.13), PM: 1174 (0.10), AM: 615 (0.13), A: 62 (0.16) |
| 2 | 5 | 3810 (0.11) | PM: 570 (0.14), AM: 519 (0.13), A: 272 (0.10) |
| 3 | 1 | 2821 (0.16) | V1: 2775 (0.16), PM: 14 (0.14), RL: 26 (0.12) |
| 3 | 2 | 2681 (0.09) | V1: 2092 (0.09), PM: 549 (0.10) |
| 3 | 3 | 3432 (0.22) | V1: 2665 (0.21), RL: 228 (0.19), AL: 159 (0.20), LM: 346 (0.34) |
| 3 | 4 | 3931 (0.14) | V1: 2399 (0.16), PM: 1425 (0.12), AM: 66 (0.17) |
| 3 | 5 | 3060 (0.23) | V1: 2004 (0.21), RL: 1 (0.00), AL: 118 (0.20), LM: 773 (0.29) |
| 3 | 6 | 3760 (0.22) | V1: 1222 (0.26), PM: 1636 (0.21), AM: 479 (0.17), A: 341 (0.15) |
| 4 | 1 | 3071 (0.26) | V1: 1280 (0.34), A: 15 (0.20), RL: 1150 (0.22), AL: 288 (0.27), LM: 98 (0.24) |
| 4 | 2 | 3718 (0.20) | V1: 1980 (0.20), PM: 361 (0.25), AM: 326 (0.21), A: 722 (0.19), RL: 264 (0.18) |
| 4 | 3 | 3676 (0.29) | V1: 1038 (0.34), RL: 1276 (0.29), AL: 616 (0.32), LM: 238 (0.34) |
| 4 | 4 | 3998 (0.12) | V1: 1825 (0.13), PM: 853 (0.11), AM: 717 (0.11), A: 496 (0.09), RL: 29 (0.21) |
| 4 | 5 | 3727 (0.30) | V1: 1914 (0.35), A: 77 (0.04), RL: 1308 (0.25), AL: 159 (0.40), LM: 53 (0.38) |
| 4 | 6 | 3283 (0.24) | V1: 971 (0.33), A: 337 (0.20), RL: 1189 (0.24), AL: 78 (0.28) |
| 5 | 1 | 3756 (0.19) | V1: 1036 (0.16), PM: 1017 (0.18), AM: 1158 (0.20), A: 487 (0.24) |
| 5 | 2 | 3392 (0.22) | V1: 2514 (0.22), PM: 270 (0.18), AM: 174 (0.23), A: 198 (0.18), RL: 198 (0.31) |
| 5 | 3 | 2888 (0.33) | V1: 1126 (0.43), PM: 5 (0.40), AM: 2 (0.00), A: 770 (0.23), RL: 493 (0.37) |
| 5 | 4 | 3136 (0.27) | V1: 1500 (0.32), PM: 210 (0.23), AM: 143 (0.23), A: 896 (0.21), RL: 249 (0.34) |
| 5 | 5 | 3161 (0.23) | V1: 1165 (0.28), PM: 644 (0.20), AM: 806 (0.21), A: 494 (0.18), RL: 3 (0.33) |
| 5 | 6 | 2876 (0.18) | V1: 991 (0.20), PM: 697 (0.15), AM: 770 (0.16), A: 358 (0.18) |
| 6 | 1 | 3441 (0.20) | V1: 1593 (0.21), PM: 193 (0.20), AM: 110 (0.21), A: 862 (0.22), RL: 496 (0.19) |
| 6 | 2 | 3575 (0.29) | V1: 2681 (0.28), RL: 214 (0.36), AL: 222 (0.31), LM: 417 (0.35) |
| 6 | 3 | 3757 (0.18) | V1: 2239 (0.21), PM: 1256 (0.14), AM: 227 (0.14) |
| 6 | 4 | 2479 (0.25) | V1: 1789 (0.25), PM: 26 (0.19), AM: 8 (0.13), A: 192 (0.21), RL: 421 (0.27) |
| 6 | 5 | 2383 (0.18) | V1: 1596 (0.17), PM: 96 (0.15), AM: 55 (0.11), A: 344 (0.15), RL: 245 (0.26) |
| 7 | 1 | 2554 (0.12) | V1: 57 (0.16), PM: 570 (0.12), AM: 643 (0.14), A: 500 (0.15) |
| 7 | 2 | 2388 (0.17) | V1: 532 (0.16), PM: 895 (0.18), AM: 338 (0.21), A: 10 (0.00) |
| 8 | 1 | 3543 (0.28) | V1: 1877 (0.26), PM: 272 (0.22), AM: 165 (0.23), A: 573 (0.35), RL: 573 (0.31) |
| 8 | 2 | 3775 (0.28) | V1: 2418 (0.25), RL: 772 (0.33), AL: 272 (0.31), LM: 263 (0.32) |
| 8 | 3 | 3569 (0.19) | V1: 733 (0.30), PM: 1400 (0.16), AM: 426 (0.19), A: 13 (0.46) |
| 8 | 4 | 3565 (0.25) | V1: 2968 (0.24), PM: 123 (0.18), AM: 103 (0.19), A: 103 (0.24), RL: 234 (0.38) |
| 8 | 5 | 3824 (0.29) | V1: 2138 (0.27), A: 15 (0.33), RL: 1071 (0.33), AL: 257 (0.39), LM: 165 (0.26) |
| 8 | 6 | 3449 (0.21) | V1: 2677 (0.21), PM: 128 (0.12), AM: 73 (0.12), A: 86 (0.21), RL: 433 (0.28) |

**Table S2: Summary of neurons recorded by visual area, across mice; related to Figure 2.** Session counts reflect total unique sessions identifying neurons in that visual area.

| Visual Area | # Neurons | Response (%) (% responsive) | Total # Sessions |
|---|---|---|---|
| V1 | 63041 | 14439 (0.23) | 40 (8 mice) |
| PM | 17272 | 2780 (0.16) | 31 (8 mice) |
| AM | 11050 | 1949 (0.18) | 29 (8 mice) |
| A | 14109 | 2671 (0.19) | 32 (8 mice) |
| RL | 13885 | 3804 (0.27) | 28 (7 mice |
| AL | 2352 | 728 (0.31) | 10 (5 mice) |
| LM | 2399 | 758 (0.32) | 9 (5 mice) |

**Table S3: Summary of mice contributing neural data; related to Figure 2.**

| Mouse ID | Experimental ID | Sex | DOB | First session | Last session | # Sessions |
|---|---|---|---|---|---|---|
| 1 | 4512_1L | F | 2024-07-23 | 2025-04-23 | 2025-05-27 | 6 |
| 2 | 4514 | F | 2024-08-14 | 2025-04-23 | 2025-05-23 | 5 |
| 3 | 4518 | M | 2024-09-06 | 2025-04-23 | 2025-05-23 | 6 |
| 4 | 18870 | M | 2024-06-23 | 2025-04-23 | 2025-05-23 | 6 |
| 5 | 4575 | F | 2024-11-05 | 2025-07-16 | 2025-09-04 | 6 |
| 6 | 4576_2R | M | 2024-11-05 | 2025-07-15 | 2025-09-03 | 6 |
| 7 | 4576_2L | M | 2024-11-05 | 2025-07-15 | 2025-07-17 | 2 |
| 8 | 4611 | F | 2024-11-25 | 2025-07-16 | 2025-09-04 | 6 |

**Table S4: Summary of mice contributing behavioral data; related to Figure 1.**

| Mouse ID | Experimental ID | Sex | DOB | First Session | Last Session | # Sessions |
|---|---|---|---|---|---|---|
| B1 | 4512_1R | F | 2024-07-23 | 2024-12-17 | 2025-01-21 | 5 |
| B2 | 4512_1L | F | 2024-07-23 | 2025-03-13 | 2025-03-21 | 4 |
| B3 | 4514 | F | 2024-08-14 | 2025-03-10 | 2025-03-21 | 4 |
| B4 | 4518 | M | 2024-09-06 | 2025-03-08 | 2025-03-21 | 4 |
| B5 | 18870 | M | 2024-06-23 | 2025-03-08 | 2025-03-21 | 5 |
